# Discoidin domain receptor 2 drives melanoma drug resistance through AXL-dependent phenotype switching

**DOI:** 10.1101/857904

**Authors:** Margaux Sala, Nathalie Allain, Arnaud Jabouille, Elodie Henriet, Aya Abou-Ammoud, Arnaud Uguen, Sylvaine Di-Tommaso, Cyril Dhourte, Anne-Aurélie Raymond, Jean-William Dupuy, Emilie Gerard, Nathalie Dugot-Senant, Benoit Rousseau, Jean-Phillipe Merlio, Anne Pham-Ledart, Béatrice Vergier, Violaine Moreau, Frédéric Saltel

**Affiliations:** Inserm, UMR1053, BaRITOn, Bordeaux Research in Translational Oncology, Team 1, 146 Rue Léo Saignat, Bordeaux, F-33076, France; Bordeaux University, 146 rue Léo Saignat, Bordeaux, F-33076, France; Brest University Hospital Center, Department of Pathology, Brest, F-29200, France; Oncoprot, UMS 005, Bordeaux, F-33076, France; Bordeaux University, Plateforme Protéome, Centre for Functional Genomics, F-33000 Bordeaux, France; Bordeaux University Hospital, Dermatology Department, 1 avenue Jean Burguet, F-33000 Bordeaux, France; Histology Plateforme, UMS 005, Bordeaux, F-33076, France; Common Animal Housing Service, Bordeaux University, F-33076; INSERM, UMR1053, BaRITOn, Bordeaux Research in Translational Oncology, Team 3, “Cutaneous Lymphoma Oncogenesis”, 146 Rue Léo Saignat Bordeaux, F-33076, France; Bordeaux University Hospital, Tumor Bank and Tumor Biology Laboratory, Avenue de Magellan, Pessac, F-33604, France

**Author notes:** Ewald’s lab-dpt of Cell biology, John Hopkins University School of Medicine, Baltimore, USA. MFP, UMR 5234, Bordeaux University, 146 rue Léo Saignat, Bordeaux, F-33076, France. **Contact information**, Frédéric Saltel, Ph.D. INSERM, UMR1053, BaRITOn Bordeaux Research in Translational Oncology, 146 rue Leo Saignat, 33076 Bordeaux, France.

**Keywords:** melanoma, discoidin domain receptors, phenotype switching, resistance, vemurafenib, dasatinib

## Abstract

Anti-BRAF plus anti-MEK are used as first-line treatment of patients with metastatic melanomas harboring *BRAF* V600E mutation. The main issue with targeted therapy is acquired cellular resistance. In 70% of acquired resistance, melanoma cells switch their phenotype and become more aggressive and invasive. The molecular signature of this phenotype is MITF low, AXL high associated with actin cytoskeleton reorganization. After this switch, resistant cells present an anarchic cell proliferation due to MAP kinase pathway hyper-activation. We demonstrate that resistant cell lines presenting phenotype switching overexpress DDR1 and DDR2. We show that DDR2 inhibition induces a decrease in AXL and reduces actin stress fiber formation. Once this phenotype switching is acquired, we report that both DDRs promotes tumor cell proliferation, but only DDR2 can over-activate the MAP kinase pathway in resistant invasive cells *in vitro* and *in vivo*. Therefore, DDRs inhibition could be a promising strategy for countering this resistance mechanism.

**Significance:** Our results show that DDR2 is a relevant target in melanoma resistance. DDR2 is required at the beginning of resistance for melanoma cell phenotype switching to occur. After phenotype switching, DDRs promote tumor cell proliferation of resistant invasive melanoma cells, but only DDR2 is able to over-activate the MAP kinase pathway. We put forward dasatinib (a DDR inhibitor) as a potential second-line treatment after targeted dual therapy for resistant patients overexpressing DDRs.

## Introduction

Melanoma, a malignant transformation of melanocytes, is the most aggressive form of skin cancer. In 2018, approximately 300 000 new melanoma cases were diagnosed and 60 000 deaths worldwide^1^. Before 2011, overall patient survival was six months. However, due to the development of combined targeted therapy and immunotherapy, overall survival is now up to two years^2^.

The most common type of cutaneous melanoma is Superficial Spreading Melanoma (SSM), accounting for 70% of all melanomas^3^. The *BRAF* V600E mutation is observed in almost 60% of SSM^4^. The *BRAF* gene encodes a protein kinase involved in the MAP kinase pathway. Its mutation induces a constitutive activation of BRAF, leading to over-activation of downstream signaling pathways, notably MEK and ERK. This in turn promotes anarchic cell proliferation and cell invasion in melanoma^5^. Over the past few years, new therapies have been developed, including targeted bi-therapy and more recently immunotherapy. However, the response rate to immunotherapy is only 15% and a severe toxicity is observed in more than 25% of patients^2^. The combination of two treatments, an anti-BRAF plus an anti-MEK (vemurafenib and cobimetinib, or dabrafenib and trametinib), is still currently used as first-line treatment in the management of patients with metastatic melanomas harboring the somatic *BRAF* V600E mutation^6,7^. Vemurafenib and dabrafenib inhibit the activity of the mutant BRAF protein, whereas trametinib and cobimetinib inhibit MEK protein expression^8–10^. However, the main problem related to targeted therapy is the acquired cellular resistance found in 80% of patients approximately two years after treatment starts^11^. In 70% of acquired resistance to targeted therapy, melanoma cells tend to change their molecular and cellular phenotypes in an epithelial-to-mesenchymal transition (EMT) called phenotype switching^12^. Melanoma cells switch into an invasive state and therefore become much more aggressive. The phenotype switching signature is characterized by AXL high, MITF low, and actin cytoskeleton remodeling^13–15^. Several studies have demonstrated an important role for AXL in melanoma given its expression level is often elevated and inversely correlated with MITF expression patterns in BRAF inhibitor-resistant melanomas^12,14,16^. MITF is a transcription factor involved in lineage-specific pathway regulation of melanocytes, and AXL is a tyrosine kinase receptor involved in resistance to therapy in different cancers, including melanoma^17^,. Despite targeted bi-therapy treatment, after phenotype switching resistant melanoma cells present an anarchic cell proliferation due to hyper-activation of MAP kinase activity^11^. This hyper-activation could be due to genetic events, including *NRAS* or *BRAF* mutations, *BRAF* splice variants, secondary mutation of *MEK*, overexpression of the BRAF oncoprotein, or upregulation of tyrosine kinase receptors, such as PDGFR, EGFR, or FGFR^13,18–21^. Therefore, in this study we searched for deregulated receptors after vemurafenib treatment that could participate to this phenotype switching, and broadly to the resistance. We highlight the deregulation of DDRs and focus on their involvement in melanoma drug resistance.

DDRs are composed of two members: DDR1 and DDR2^22^. These transmembrane receptors are activated by collagens in their native triple-helix form^23–25^. DDR1 is activated for instance by type I and IV collagens, whereas DDR2 preferentially binds type I, II, and X collagens^23,25^. Moreover, DDRs are involved in several physiological functions and have been found overexpressed in a large number of cancer subtypes. where they are associated with cell proliferation, invasion, migration, and drug resistance^26,27^. Previous reports have indicated that DDRs can activate the MAP kinase pathway^26,28–31^. DDR1 is involved in cancer therapy resistance by activating Cox2 expression through NFκB pathways in breast cancer cells^32^. DDR1 also promotes tumor cell resistance in lymphoma, ovarian cancer, and glioblastoma cells^33^. Overexpression of DDR2 is associated with breast cancer recurrence through activation of the Erk/Snail 2 pathway^33^. In a recent study, DDR2 was shown to activate the Src-PI3K/Akt-NFκB signaling pathway, allowing the expression of apoptosis-inhibiting proteins^34^. Most studies on DDRs only focus on one of the two receptors; none of them integrate the role of both DDRs in melanoma, whereas both receptors are expressed. Moreover, no study has yet investigated the involvement of DDRs in the resistance to targeted therapies in melanoma. Therefore, the aim of this study was to fully investigate the shared and distinct roles of DDRs in resistance to targeted therapies in melanoma in order to determine their potential as new targets for countering the resistance mechanism.

Altogether, our data uncover that DDR2 is specifically involved in phenotype switching of resistant cells through AXL regulation and RhoA activation. We additionally show that both DDRs promote cell proliferation after phenotype switching, but only DDR2 is able to overactivate the MAP kinase pathway. Our study highlights that targeting these receptors, and especially DDR2, could reduce tumor progression. Moreover, we put forward dasatinib, an FDA-approved drug targeting DDRs, as a second-line treatment after targeted bi-therapy for resistant patients overexpressing DDRs.

## Materials and methods

### Cell culture

Human melanoma cell lines with *BRAF* V600E mutation (229, 238 and 249) have been previously described^20^. These cell lines were a generous gift from Dr Sophie Tartare-Deckert (U1065, Nice). The resistant cells (named 229 R, 238 R and 249R) were derived from the sensitive cells (named 229 S and 238 S) following vemurafenib treatment^35^. Both sensitive and resistant cell lines, 229 S/R, 238 S/R, 249 S/R were cultured in Dulbecco’s modified Eagle’s medium with 4.5 g/l glucose Glutamax-I (Invitrogen) supplemented with 10% fetal calf serum (Sigma-Aldrich). The resistance was maintained by addition of 1 μM vemurafenib (LC Laboratories) to the medium. Vemurafenib was added every day throughout experiments with resistant cells.

### Collagen I coating

Collagen polymerization was carried out as described previously^36^. In brief, collagen was diluted to 0.5 mg/ml in DPBS 1X, then polymerized for 4 h at 37 °C before cell seeding. Cells were seeded for 4 h on collagen before fixation.

### Immunofluorescence

Cells were fixed with 4% paraformaldehyde (pH 7.2) for 10 min, permeabilized with 0.2% Triton X-100 for 10 min, and incubated with various antibodies. Cell imaging was performed with an SP5 confocal microscope (Leica) using a 63×/NA 1.4 Plan Neofluor objective lens. Each channel was imaged sequentially using the multitrack recording module before merging in order to prevent contamination between fluorochromes.

### Reagents and drugs

Vemurafenib and dasatinib were purchased from LC laboratories. The DDR2 inhibitor (CR-13452) was a generous gift from Prof. Gregory Longmore’s laboratory (ICEE institute, St Louis). The DDR1 inhibitor (7rh) was purchased from Sigma.

### Western Blots

Protein was extracted from cell lysates using the radio-immunoprecipitation assay buffer (25 mM Tris HCl, pH 7.5, 150 mM NaCl, 1% IGEPAL, 1% sodium deoxycholate, and 0.1% SDS) completed with protease and phosphatase inhibitors. Proteins were blotted on a nitrocellulose membrane (Transblot^®^ TurboTM midi-size, Bio-Rad), blocked with the Odyssey blocking buffer (LI-COR Company), and probed with primary antibodies overnight at 4°C. The following antibodies were used: rabbit monoclonal anti-DDR1 (5583S, Cell Signaling), rabbit monoclonal anti-DDR2 (12133S, Cell Signaling), rabbit monoclonal anti-P-DDR1 (14531, Cell Signaling), rabbit monoclonal anti-P-DDR2 (MAB25382, R&D Systems), mouse monoclonal anti-P-Erk (9106S, Cell Signaling); rabbit monoclonal anti-Erk (9102S, Cell Signaling); rabbit monoclonal anti-P-AKt (2965S, Cell Signaling), rabbit monoclonal anti-Akt (9272S, Cell Signaling), anti-AXL (8661S, Cell Signaling), anti-MITF (sc-56725, Santa Cruz) and mouse monoclonal anti-GAPDH (sc-166545, Santa Cruz Biotechnology). Primary antibodies were diluted in Tris-buffered saline (TBS) with 5% bovine serum albumin (BSA). After washing with TBS 0.1% Tween (twice for 10 minutes), membranes were incubated for 1 h with the corresponding species-specific fluorescent far-red coupled secondary antibody: IRDye 680RD goat (polyclonal) anti-rabbit IgG (H+L) (LI-COR) or IRDye 800RD goat (polyclonal) antimouse IgG (H+L) (LI-COR) diluted 1:5000 in TBS 5% BSA. After washing twice for 10 minutes with TBS 0.1% Tween and with 1x TBS, membranes were revealed with the BioRad imager with the Image Studio software as recommended by the manufacturer. Quantification of the correct size band for each antibody was performed with the ImageLab software.

### Transfections

Small interfering RNAs (siRNA) (20 nM) were transfected using Lipofectamine RNAiMax (Invitrogen) according to the manufacturer’s instructions. The siRNA sequences targeting human DDR1 and DDR2 were as follows: siDDR1 5’–GAAUGUCGCUUCCGGCGUGUU-3’ and siDDR2 5’–GAAACUGUUUAGUGGGUAA-3’. A control siRNA targeting luciferase (CT) 5’–CGTACGCGGAATACTTCGA-3’ was purchased from Eurofins MWG Operon, Inc. The efficiency of the silencing was determined using western blotting and reverse transcription-quantitative polymerase chain reaction (RT-qPCR).

### RT-qPCR

mRNAs were extracted from cultured cells using the Nucleospin RNA^®^ kit from Macherey Nagel according to the manufacturer’s instructions. cDNA was synthetized from 1 μg of total RNA with Maxima Reverse Transcriptase (Fermentas). 30 ng of cDNA were then subjected to PCR amplification on an RT-qPCR system using the CFX96 Real Time PCR detection system (Biorad). The SYBR^®^ Green SuperMix for iQTM (Quanta Biosciences, Inc.) was used with the following PCR amplification cycles: initial denaturation (95 °C for 10 min), followed by 40 cycles of denaturation (95 °C for 15 s) and extension (60 °C for 1 min). Gene expression results were first normalized to an internal control with 18S ribosomal RNA. Relative expression levels were calculated using the comparative (2-ΔΔCT) method. All primers used in RT-qPCR experiments are listed in supplementary Table 1.

### IncuCyte® assays

**Proliferation assays:** Cells were seeded in 96-well plates (5 000 cells per well) and then monitored by the IncuCyte® Zoom videomicroscope (Essen Bioscience). Cell confluence and quantification was measured using the IncuCyte® imaging system. For dasatinib experiments, dasatinib was added at 100 nM. **Apoptosis assays:** 229 R and 238 R cells were seeded at a density of 5 000 cells per well and left to grow overnight. The next day, dasatinib (100 nM) was prepared and added, directly followed by addition of the caspase-3/7 reagent at a final dilution of 1/1000. Cells were then incubated in the IncuCyte® Zoom live cell imaging system. The green reagent from the IncuCyte® Caspase-3/7 Apoptosis Assay couples the activated caspase-3/7 recognition motif (DEVD) to NucView™ 488, a DNA intercalating dye, in order to enable quantification of apoptosis over time. **Spheroid assays:** Cells were seeded into 96-well ultra-low attachment plates. Then, we monitored spheroid formation using IncuCyte^®^ videomicroscopy for 72 h. After spheroid formation, treatment (dasatinib or CR-13452) was added. The quantifications of spheroid area and perimeter were performed using Image J.

### Quantification G-actin/F-actin

The G-actin/F-actin ratio in 229 R cells was assayed using the G-actin/F-actin In Vivo Assay Kit (Cytoskeleton, Denver, CO, USA). 229 R cells were scraped in LAS2 buffer and lysed. Then, lysates were centrifuged to produce an unbroken cell pellet. We centrifuged the supernatant fraction at 100 000 g to separate F-actin from soluble G-actin. The supernatant and the pellet were then analyzed by immunoblotting with the anti-actin antibody provided in the kit.

### Immunoprecipitation

Three mg/500 μL of solubilized proteins were incubated with mCherry antibody overnight at 4°C. Fifty μL of protein G-conjugated sepharose beads were added for a 1-h incubation at 4°C. After 5 washes in lysis buffer, the pellet was resuspended in Laemli buffer for gel electrophoresis.

### Invasion assay

Transwell invasion assays were performed in 24-well plates with 8-μM BioCoat control inserts. A total of 5*10^4^ cells ((229 S, D-DDR2 mCh, R), (238 S, S-DDR2 mCh, R) and (249 S, D-DDR2 mCh, R)) were prepared in DMEM. 150 μL cell suspensions were added into the upper 24-well transwell chamber with Matrigel (1 mg/mL). After 24 hours, cotton swabs were used to remove the non-invasive cells from the upper chambers, and the cells in the lower chambers were fixed with crystal violet (Sigma-Aldrich) for 20 minutes.

### Proteomic analysis

Cell lysis was performed in RIPA Buffer. The sample preparation and protein digestion steps were performed as previously described^37^. NanoLC-MS/MS analyzes were performed using an Ultimate 3000 RSLC Nano-UPHLC system (Thermo Scientific, USA) coupled to a nanospray Orbitrap Fusion™ Lumos™ Tribrid™ mass spectrometer (Thermo Fisher Scientific, California, USA). The Mascot 2.5 software was used for protein identification in batch mode by searching against the UniProt Homo sapiens database (74 489 entries, Reference Proteome Set, release date: May 16, 2019) from http://www.uniprot.org/. Raw LC-MS/MS data were imported into Proline Studio (http://proline.profiproteomics.fr/) for feature detection, alignment, and quantification. A protein identification was only accepted if it had at least two specific peptides with a pretty rank=1 and a protein FDR value below 1.0% calculated using the “decoy” option in Mascot. Label-free quantification of MS1 spectra by extracted ion chromatograms (XIC) was carried out with the parameters previously indicated ^37^. Protein ratios were median normalized.

Gene Set Enrichment Analysis (GSEA) was performed against the Ingenuity Pathways Database (Canonical Pathways). Only shared and significantly deregulated pathways were considered.

### Xenograft mouse model

The institutional animal ethics committee of Bordeaux University approved all animal use procedures and all efforts were made to minimize animal suffering. Five million 229 R cells were resuspended in a mix 1: 1 with DMEM 4.5 g/l glucose Glutamax-I medium and Matrigel. The mixture was then injected subcutaneously into the right flanks of anesthetized 8-week-old NOD/LtSz-*scid* IL2Rγ*null* (NSG) mice. Tumor formation and tumor volume, based on caliper measurements, were monitored twice a week. The mice were treated with vemurafenib until the tumors reached approximately 150 mm^3^ in volume. Subsequently, the mice were randomly assigned into two groups: a control group of mice with ongoing treatment with vemurafenib (40 mg/kg) by oral gavage, and a second group of mice treated with dasatinib (20 mg/kg) by oral gavage (n=5 for each treatment).

### Immunohistochemistry

Primary tumors were fixed in 10% buffered formaldehyde. Selected representative tissue slides for tumors treated with or without dasatinib were processed for Annexin V immunohistochemistry (IHC). Tissue sections (2.5 μm thick) were dewaxed and rehydrated, and antigens were retrieved in sodium citrate buffer (pH6) for 20 mins. All staining procedures were performed in an Autostainer (Dako-Agilent Clara, United States) using standard reagents provided by the manufacturer. The sections were incubated with an anti-Annexin V (Abcam, ab14196) rabbit polyclonal antibody at a 1μg/ml dilution for 45 min at room temperature. EnVision Flex/HRP (Horseradish peroxidase) (Dako-Agilent) was used for signal amplification for 20 minutes. 3,3’-Diamino-benzidine (DAB, Dako) development was used for detecting primary antibodies. The slides were counterstained with hematoxylin, dehydrated, and mounted. Each immunohistochemical run contained a negative control (buffer replacing the primary antibody). Sections were visualized with a Nikon-Eclipse501 microscope and images were acquired using NIS-Elements F.

### Statistical Analysis

Data were reported as the mean ± SEM of at least three experiments. Statistical significance (P < 0.05 or less) was determined using a paired *t* test or analysis of variance (ANOVA) as appropriate and performed with GraphPad Prism software (GraphPad Software). Significance levels were as follows: * p<0.05, ** p≤0.01, *** p≤0.001.

## Results

### Overview of DDR expression in melanoma progression

To investigate the expression of DDRs in melanoma, we compared their protein expression level in three benign nevi versus a series of 30 metastatic patient biopsies and 14 melanoma cell lines. DDRs were poorly expressed in nevi, whereas DDR1, DDR2, or both were found expressed in 77% of the metastatic biopsies analyzed. Furthermore, we showed that DDRs were expressed in 78.6% of the 14 different metastatic melanoma cell lines tested (**Figure 1A, supplemental 1A & 1B)**. These results show that DDR1 and/or DDR2 expression at the protein level are associated with metastatic melanoma. TCGA database analysis on cutaneous melanoma demonstrated that only an increase in DDR2 mRNA expression level was associated with metastasis and thus to bad prognosis (**Figure 1B**). Among treated patients with melanoma, 80% will develop a resistance to anti-BRAF targeted therapy^11^. Accordingly, we further analyzed DDR expression before and after an anti-BRAF treatment. To do so, we explored RNA sequencing data (*GSE50509*) from eight patients before and after vemurafenib treatment. We found overexpressed DDR1 in 50% of the cases following vemurafenib treatment, while DDR2 was overexpressed in 62.5% of the cases. Moreover, we observed an overexpression of DDR1, DDR2, or both in 75% of the cases (**Figure 1C)**. We obtained the same results from the analysis of an RNA sequencing database from 16 patients before and after treatment with dabrafenib (another anti-BRAF). Following dabrafenib treatment, we observed a total of 87.5% of all cases exhibiting a DDR overexpression (**Supplemental 1C**).

**Figure 1:**
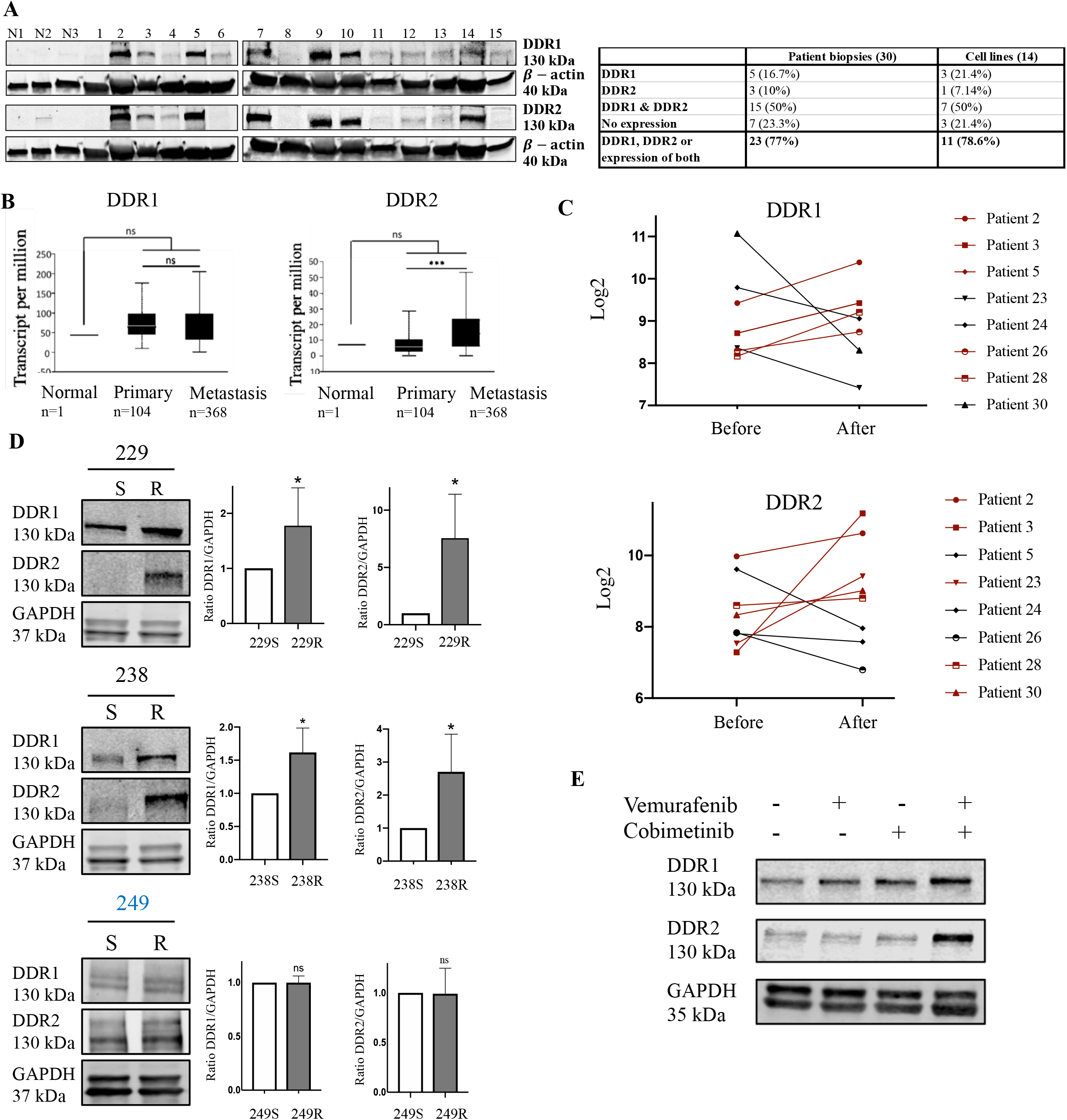
Overview of DDR expression in melanoma progression. **A**) Western blot analysis of DDR1 and DDR2 expression in 3 benign nevi (N1, N2, N3), 15 metastatic melanoma patient biopsies (1–15), and in 14 melanoma cell lines. β-actin was used as the endogenous loading control. **B**) DDR expression in melanoma progression from a metaanalysis of 363 cutaneous melanomas from TCGA database analysis (skin cutaneous melanoma, PanCancer Atlas. **C**) RNA sequencing data (*GSE50535, GSE5050*) for DDR1 or DDR2 mRNA expression before and after vemurafenib treatment. **D**) Western blot analysis of DDR1 and DDR2 expression in a subset of three melanoma cell lines (229, 238, 249) either sensitive (S) or resistant (R) to vemurafenib. GAPDH was used as the endogenous loading control. The graph shows the quantification of DDR expression. Values are expressed as the mean ± SEM of three independent experiments. *p<0.05, ns: non-significant. **E**) Western blot analysis of DDR1 and DDR2 expression in 229 S cells treated with vemurafenib (10 nM), cobimetinib (10 nM), or both over 2 months. GAPDH was used as the endogenous loading control.

Subsequently, in order to confirm our results we analyzed DDR expression at mRNA and protein levels in three isogenic pairs of melanoma cell lines: 229, 238, and 249 of parental sensitive (S) and resistant (R) cells to vemurafenib^35^. The resistant melanoma cell lines were derived from the sensitive *B-RAF*-mutated cell lines by repeated vemurafenib treatment^35^. At the protein expression level, we demonstrated that DDR1 and DDR2 were both overexpressed in 229 R and 238 R cells compared to sensitive cells. Precisely, we observed a 1.5-fold increase in DDR1 expression in both resistant cell lines, and a 2-fold increase in 238 R cells and a 6-fold increase in 229 R cells in DDR2 expression. There was no change in DDR1 or DDR2 expression in 249 R cells (**Figure 1D**).

Due to the fact that patients with melanoma are currently treated in the clinical setting with bi-therapy, we examined DDR expression in sensitive melanoma cells treated with vemurafenib, cobimetinib, or both over two months. We demonstrated that 229 S cells treated with bi-therapy exhibited a DDR overexpression, similarly to vemurafenib resistant cells (**Figure 1E**). These data demonstrate that drug-resistant melanoma cells overexpress DDR1 and DDR2.

### DDR2 involvement in AXL expression

It is known that 229 R and 238 R cells are in an invasive state (MITF low, SOX10 low, AXL high) and have undergone phenotype switching^14,20,38^. Conversely, 249 R cells are in a proliferative state (MITF high, SOX10 high, AXL low). These findings prompted us to explore a possible link between DDRs and phenotype switching.

We observed that AXL, a marker of invasion, and DDR2 (but not DDR1) were overexpressed at the mRNA level in resistant invasive melanoma cells (**Figure 2A**). Analysis of an additional RNA sequencing database confirmed that DDR2 (but not DDR1) and AXL were overexpressed in invasive melanoma cells (**Supplemental 2A**). At protein level, we demonstrated that DDR1 and DDR2 were both overexpressed in resistant cells with an invasive phenotype compared to sensitive cells. DDR overexpression at the protein level was correlated with an overexpression of AXL (**Figure 2B**). These results suggest that DDR expression at the protein level is linked to the resistant invasive phenotype.

**Figure 2:**
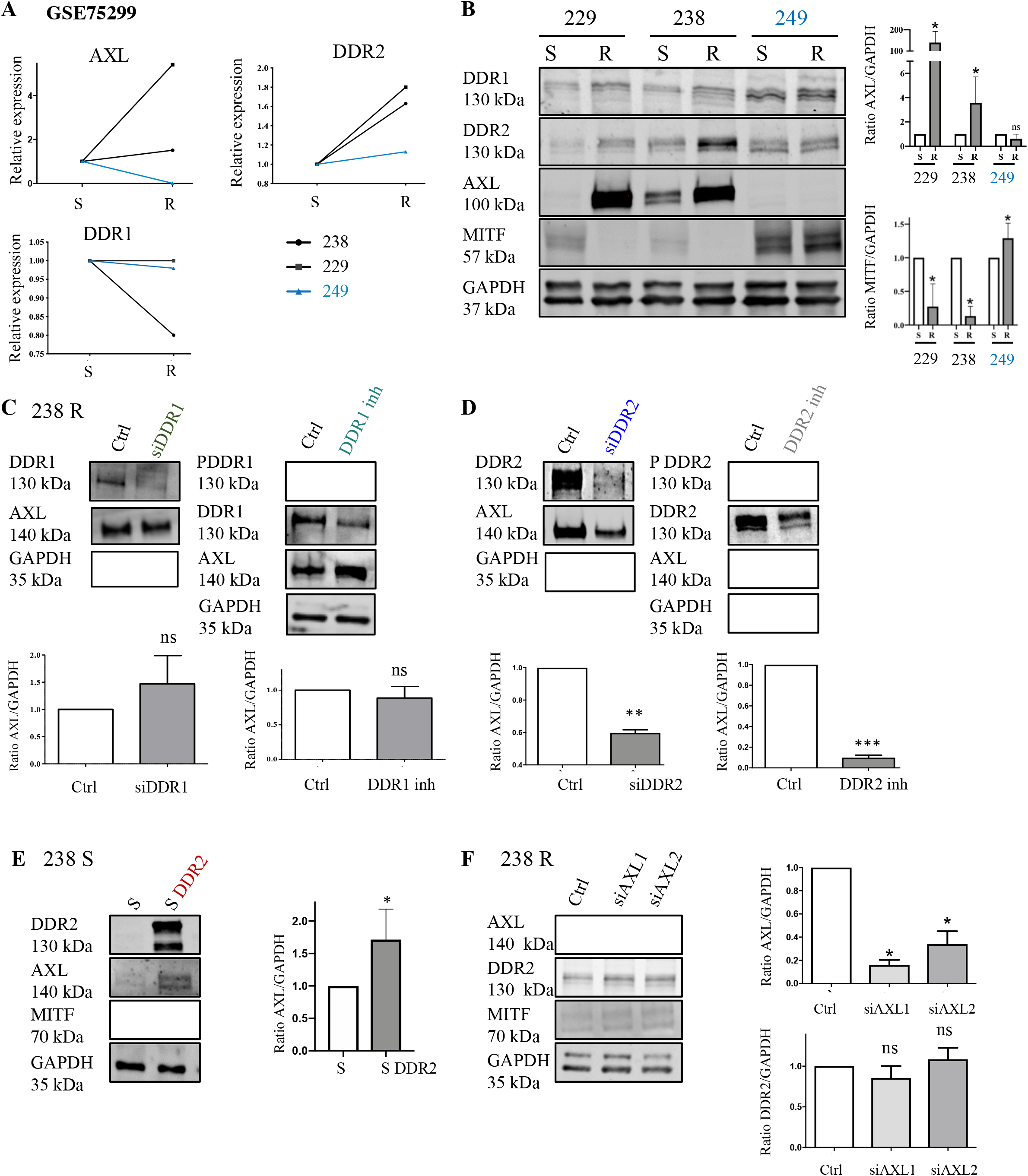
DDR2 regulates AXL expression in resistant invasive melanoma cells. **A**) RNA sequencing data (*GSE65185*) for AXL, DDR1 and DDR2 mRNA expression in vemurafenib sensitive or resistant cell lines. **B**) Western blot analysis of AXL, MITF, DDR1, and DDR2 expression in a subset of three melanoma cell lines (229, 238, 249) sensitive (S) or resistant (R) to vemurafenib. GAPDH was used as the endogenous loading control. The graph shows the quantification of AXL and MITF expression. Values are expressed as the mean ± SEM of three independent experiments. ns: non-significant, *p<0.05. **C**) 238 R cells were transfected with an siRNA control (siGl2) or targeting DDR1 (siDDR1), or 238 R cells were treated with DDR1 inhibitor. Protein extracts were then analyzed by immunoblotting to determine PDDR1, DDR1, and AXL expression. The graph shows the quantification of AXL/GAPDH expression. GAPDH was used as the endogenous loading control. Values are expressed as the mean ± SEM of three independent experiments. ns: non-significant. **D**) 238 R cells were transfected with an siRNA control (siGl2) or targeting DDR2 (siDDR2), or 238 R cells were treated with DDR2 inhibitor. Protein extracts were then analyzed by immunoblotting to determine PDDR2, DDR2, and AXL expression. GAPDH was used as the endogenous loading control. The graph shows the quantification of AXL/GAPDH expression. Values are expressed as the mean ± SEM of three independent experiments. **p<0.01, ***p<0.001. **E**) 238 S cells were transiently transfected with DDR2-mCherry. Protein extracts were then analyzed by immunoblotting to determine DDR2 and AXL expression. GAPDH was used as the endogenous loading control. The graph shows the quantification of DDR expression. Values are expressed as the mean ± SEM of three independent experiments. *p<0.05. **F**) 238 R cells were transfected with an siRNA control (siGl2) or targeting AXL (siAXL1/2). Protein extracts were then analyzed by immunoblotting to determine AXL, DDR2, and MITF expression. GAPDH was used as the endogenous loading control. Values are expressed as the mean ± SEM of three independent experiments. *p<0.05, ns: non-significant.

As mentioned before, AXL is a marker of phenotype switching. To investigate the involvement of DDRs in phenotype switching, we first determined the effects of DDRs depletion by siRNA or inactivation using specific inhibitors in resistant invasive melanoma cells. To target DDR1, we used siRNA knockdown or a specific DDR1 inhibitor (7rh inhibitor) in 238 R cells. DDR1 depletion or inhibition did not alter AXL expression (**Figure 2C**). We then analyzed the effect of DDR2 depletion by using an siRNA directed against DDR2. This time we demonstrated a decrease in AXL expression associated with DDR2 depletion (**Figure 2D**). Recently, WRG28 was identified as an allosteric inhibitor of DDR2. It is highly selective and can dissociate preformed DDR2 collagen complex and inhibit kinase-independent receptor function^39^. Based on this, we analyzed the effects of an analog of this DDR2 inhibitor (CR-13452) on AXL expression level in 238 R cells. Consistent with the result obtained with DDR2-targeting siRNA, we found that DDR2-inhibitor treatment of resistant melanoma cells was associated with a decrease in AXL expression in 238 R (**Figure 2D**) and 229 R (**Supplemental 2B)** cells. The efficacy of the inhibitor to target DDR2 is attested by loss of the phosphorylated form of DDR2. Altogether, these results demonstrate that DDR2, but not DDR1, regulates AXL expression.

In order to further confirm this result, we analyzed the effect of DDR2 overexpression on AXL expression in 238 S cells that do not undergo phenotype switching. We report an increase in AXL expression in 238 S cells overexpression DDR2 compared to sensitive control cells, and this effect did not alter MITF expression (**Figure 2E**). We also studied the effect of AXL depletion on DDR2 expression in resistant invasive cells. We showed that AXL depletion had no effect on DDR2 expression, showing that DDR2 regulates AXL and not the inverse (**Figure 2F**). Taking this further, we studied the molecular link between DDR2 and AXL. AXL is able to form heterodimers with some receptors. Therefore, we tested the potential for interaction between AXL and the DDR2 receptor. We performed AXL and DDR2 coimmunoprecipitation experiments. As shown in **Supplemental 2C**, we found DDR2 was not associated with AXL in the absence or presence of the DDR ligand of type I collagen. Indeed, DDR2 could modulate AXL expression without any interaction. In all, these results indicate that DDR2 could regulate AXL expression in melanoma cells that have undergone phenotype switching.

### The role of DDR2 in cytoskeletal remodeling during phenotype switching

In order to investigate pathways modulated by DDR2 overexpression in resistant cell lines, we used the approach of mass spectrometry on 238 R cells after transfection with DDR2-targeted siRNA or DDR2 inhibitor treatment. Using GSEA on the ratio of control/DDR2 inhibitor and siGL2/siDDR2, we noticed that RhoA and actin cytoskeleton signaling were commonly deregulated in these two conditions (**Figure 3A, supplemental 2D**). Indeed, the feature of phenotype switching corresponded to both the enrichment of actin stress fibers and a fibroblastlike morphology in 238 R and 229 R cells versus parental cell lines. On the contrary, the actin cytoskeleton of 249 R cells showed no significant differences to their sensitive parental cell lines (**Figure 3B**). These results confirmed the results of *Misek et al*. and *Girard et al*. demonstrating an actin cytoskeleton reorganization only in invasive resistant melanoma cells^40,41^. Given our mass spectrometry data showed that DDR2 modulation affects the RhoA signaling pathway, we thus studied the effect of DDR2 depletion or inhibition on actin stress fiber formation. In 229 R cells using immunofluorescence and quantification of the F-actin/G-actin ratio, we showed a decrease in actin stress fiber formation following DDR2 depletion or inhibition in resistant cells (**Figure 3C**).

**Figure 3:**
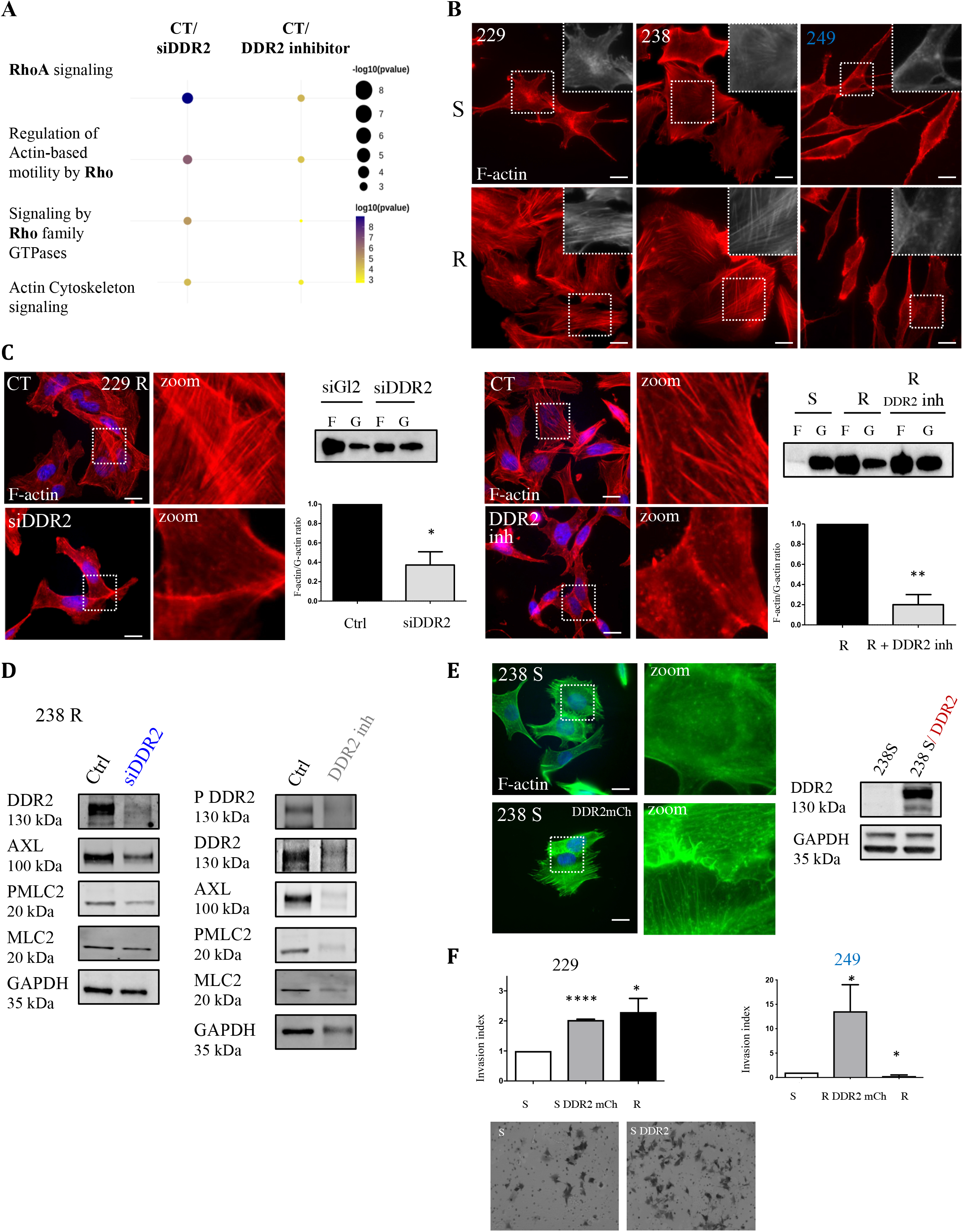
DDR2 mediates stress fiber formation in resistant invasive melanoma cells. **A)** Bubble plot of the pathways commonly and significantly enriched between these conditions: 238 R cells transfected with siRNA targeting DDR2 or treated with DDR2 inhibitor. The bubble plot represents the ratio of control/siDDR2 or control/DDR2 inhibitor. Gene set enrichment analysis (GSEA) was performed against the Ingenuity Pathways database (Fisher’s Exact test expressed in −log10pvalue). Colors and dot size represent minus logarithms of adjusted p-values (padj). Column height represents the numbers of genes enriched in a pathway. **B**) Representative images of parental versus resistant cell actin cytoskeletons. Fluorescent staining corresponds to F-actin (red) and nuclei (blue). Scale bar: 6.78 μM. **C**) Representative images of the actin cytoskeleton of resistant invasive cells transfected with siRNA targeting DDR2 or treated with DDR2 inhibitor. Fluorescent staining corresponds to F-actin (red) and nuclei (blue). Scale bar: 6.78 μM. F-actin and G-actin in each condition were separated and measured using immunoblot analysis. The graph shows the quantification of F-actin/G-actin ratio. Values are expressed as the mean ± SEM of three independent experiments, *p<0.05, **p<0.01. **D**) 238 R cells were transfected with an siRNA control (siGl2) or targeting DDR2 (siDDR2), or 238 R cells were treated with DDR2 inhibitor. Protein extracts were then analyzed by immunoblotting to determine PDDR2, DDR2, and PMLC2 expression. GAPDH was used as the endogenous loading control. **E**) 238 S cells were transiently transfected with DDR2-mCherry. Representative images of the actin cytoskeletons of 238 S or 238 S DDR2 mCh cells. Fluorescent staining corresponds to F-actin (green) and nuclei (blue). Scale bar: 6.78 μM. Protein extracts were then analyzed by immunoblotting to determine DDR2 expression. GAPDH was used as the endogenous loading control. **F**) 229 S cells or 249 S cells were transiently transfected with DDR2-mCherry. 229 (S, S DDR2-mCh, R) cells and 249 (S, R DDR2-mCh, R) were seeded in a Matrigel-coated chambers and invasion was assessed. The graph shows invasion index quantification. Values are expressed as the mean ± SEM from three independent experiments. *p = 0.0221; **p = 0.0057.

Stress fiber formation is mediated by RhoA activation. Therefore, we measured the effect of DDR2 depletion or inhibition on the phosphorylation status of MLC2, a downstream target of RhoA activation. Following DDR2 depletion or inhibition, PMLC2 decreased in 238 R cells (**Figure 3D**). Thus, DDR2 expression level controls RhoA activation that promotes actin stress fiber formation, one of the characteristics of phenotypic switching. To confirm this result, we analyzed the effect of DDR2 overexpression on stress fiber formation in 238 S cells. Indeed, we demonstrated an increase in stress fiber formation when DDR2 was overexpressed in these cells (**Figure 3E**).

Furthermore, we showed that in proliferative resistant melanoma cells (249 R), DDR2 overexpression promoted a modification in cell morphology. Accordingly, cells were more spread and exhibited an increase in stress fibers (**Supplemental 2E**). These results generally confirm our previous data indicating that DDR2 promotes stress fiber formation. Given phenotype switching is associated with an increase in cellular invasibility, we analyzed the effect of DDR2 overexpression on invasion. We showed that DDR2 overexpression enhanced cell invasion, and this was even the case for cells not having undergone phenotype switching (**Figure 3F, supplemental Figure 2F**). All these data demonstrate that DDR2 contributes to phenotype switching in melanoma cells through AXL regulation and stress fiber formation.

### DDR-associated phenotype switching is involved in MAP kinase pathway activation

After phenotype switching, resistant invasive melanoma cells will still proliferate through overactivation of the MAP kinase pathway despite targeted bi-therapy. Firstly, we confirmed that MAP kinase was over-activated in resistant invasive cells (229 R and 238 R) compared to sensitive cells; there was no overactivation of the MAP kinase pathway in 249 R cells (**Figure 4A**). It is well characterized that DDRs are able to activate the MAP kinase pathway, showing roles in the promotion of cell proliferation in several cancers, including lung cancer and breast adenocarcinoma^42^. We hypothesized that resistant invasive melanoma cell lines over-activate the MAP kinase pathway via the induction of DDRs and promote cell proliferation.

**Figure 4:**
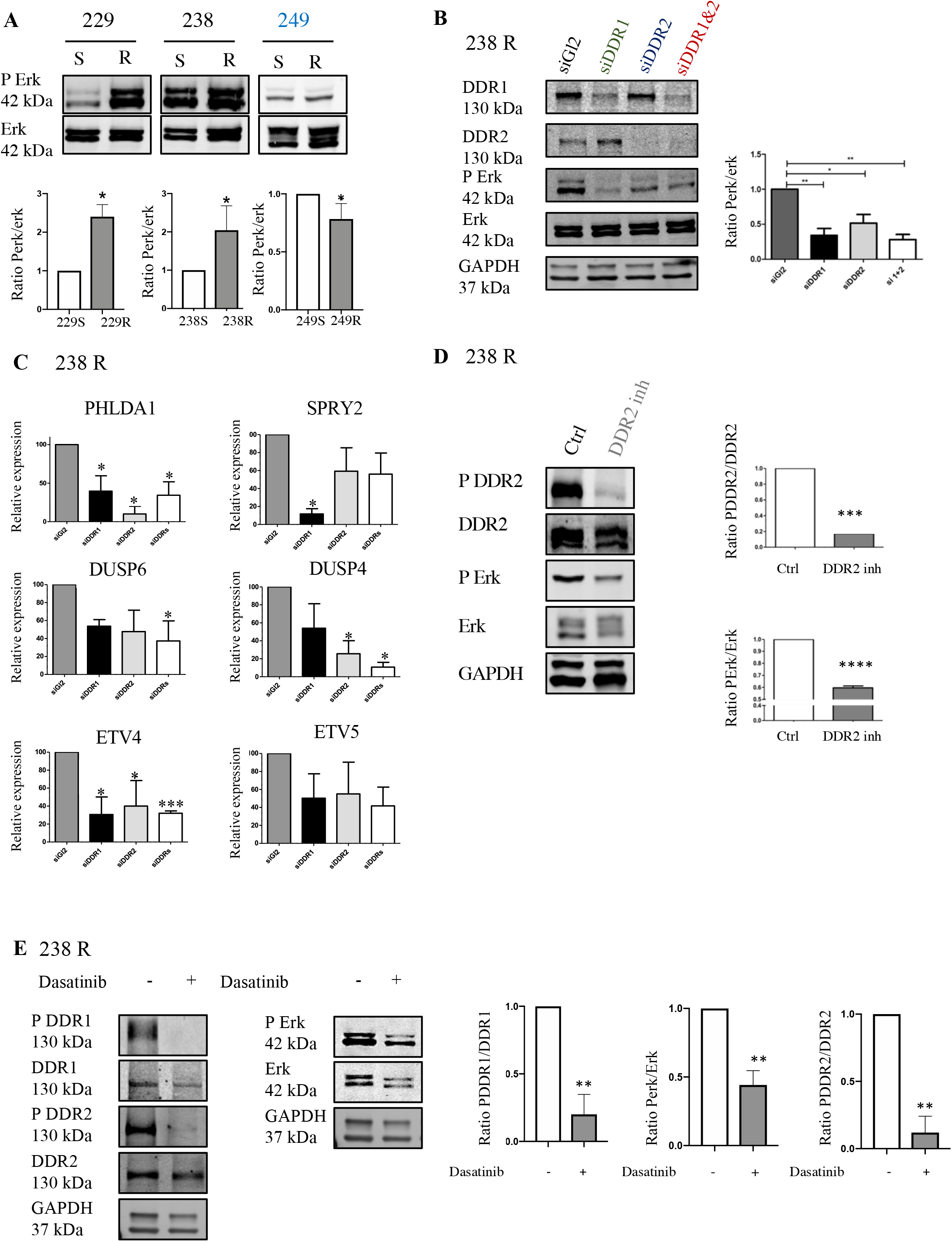
DDRs-associated phenotype switching is involved in MAP kinase pathway activation. **A**) Western blot analysis of PErk and Erk expression in a subset of three melanoma cell lines (229, 238, and 249) either sensitive (S) or resistant (R) to vemurafenib. The graph shows the quantification of PErk/Erk expression. Values are expressed as the mean ± SEM of three independent experiments *p<0.05. **B**) 238 R (left panel) cells were transfected with an siRNA control (siGl2) or targeting DDR1 (siDDR1), DDR2 (siDDR2), or both (siDDR1&2). Protein extracts were then analyzed by immunoblotting to determine PErk and Erk expression. GAPDH was used as the endogenous loading control. The graph shows the quantification of PErk/Erk expression. Values are expressed as the mean ± SEM of three independent experiments. *p<0.05, **p<0.01. **C**) mRNA expression level of MAP kinase targets in 238 R cells. The graph shows the quantification of PHLDA1, SPRY2, DUSP6, DUSP4, ETV4, and ETV5 mRNA expression level. Values are expressed as the mean ± SEM of three independent experiments. *p<0.05, ***p<0.001, ns: non-significant. **D**) 238 R cells were treated with DDR2 inhibitor (CR-13452) for 3 days. Protein extracts were then analyzed by immunoblotting to determine PErk, Erk, PDDR2, and DDR2 expression. GAPDH was used as the endogenous loading control. The graph shows the quantification of the ratio of PDDR2/DDR2 and PErk/Erk expression. **p<0.01. **E**) 238 R cells were treated with dasatinib for 2 hours. Protein extracts were then analyzed by immunoblotting to determine PErk, Erk, DDR1, and DDDR2 expression. GAPDH was used as the endogenous loading control. The graph shows the quantification of the ratio of PDDR1/DDR1, PDDR2/DDR2, and PErk/Erk expression. Values are expressed as the mean ± SEM of three independent experiments. **p<0.01.

To validate this hypothesis, we analyzed the effect of DDR depletion or DDR kinase domain inactivation on this pathway in resistant melanoma cells. We quantified the effect of DDR depletion by siRNA targeting DDR1, DDR2, or both, on MAP kinase pathway activity We found that DDR1 and/or DDR2 silencing induced a decrease in the PErk/Erk ratio in 238 R and 229 R cells (**Figure 4B, supplemental 3A**). Next, we aimed to confirm this result by studying the effect of DDR silencing on various MAP kinase targets at the mRNA expression level, including PHLDA1 and ETV4. Indeed, DDR1 and/or DDR2 depletion induced a decrease in the mRNA expression level of the majority of the MAP kinase pathway targets analyzed in both 238 R and 229 R cells (**Figure 4C, supplemental 3B**).

We analyzed the effect of DDR1 or DDR2 specific inhibitors on MAP kinase pathway activation in order to confirm our result. Surprisingly, we observed an increase in MAP kinase pathway activation using a specific DDR1 inhibitor (**Supplemental 3C**). On the other hand, we confirmed the effect of DDR2 inhibition on PErk/Erk ratio using a specific DDR2 inhibitor; this led to a decrease in MAP kinase pathway activation (**Figure 4D**). To bring this concept closer to a potential clinical use, we searched for FDA-approved drugs that inhibit kinase domains in both DDRs. We selected dasatinib, a multi-target tyrosine kinase inhibitor that inhibits DDR1 and DDR2 kinase domains simultaneously at nanomolar ranges^43^. We documented a decrease in DDR1 and DDR2 phosphorylation as expected in 238 R cells treated with 100 nM dasatinib for 2 hours. In parallel, this treatment decreased the PErk/Erk ratio in vemurafenib-resistant melanoma cells (**Figure 4E, supplemental 3D**). Altogether, these data demonstrate that both DDRs are involved in MAP kinase pathway over-activation in resistant melanoma cell lines. In addition, a DDR2 inhibitor, CR-13452 as well as dasatinib appear effective in decreasing MAP kinase activity.

### DDRs are involved in resistant tumor cell proliferation

Given DDRs play a role in MAP kinase signaling pathway activation, we investigated the biological impact of DDR silencing or inhibition on tumor cell proliferation. To do this, we analyzed cell proliferation using IncuCyte^®^ real-time cell assay monitoring. Firstly, in 2D culture we demonstrated a significant decrease in tumor cell proliferation in both resistant invasive cell lines (229 R and 238 R) when DDR1, DDR2, or both were depleted using siRNA. DDR depletion was also associated with an increase in the number of apoptotic cells, especially in conditions of double inhibition (**Figure 5A, supplemental 4A**). Therefore, we would like to study if there is an impact of DDRs depletion in proliferative melanoma resistant cell lines (249R). We demonstrated here that when DDRs are depleted in proliferative melanoma resistant cells, there is an increase of their proliferation (**Figure 5B**).

**Figure 5:**
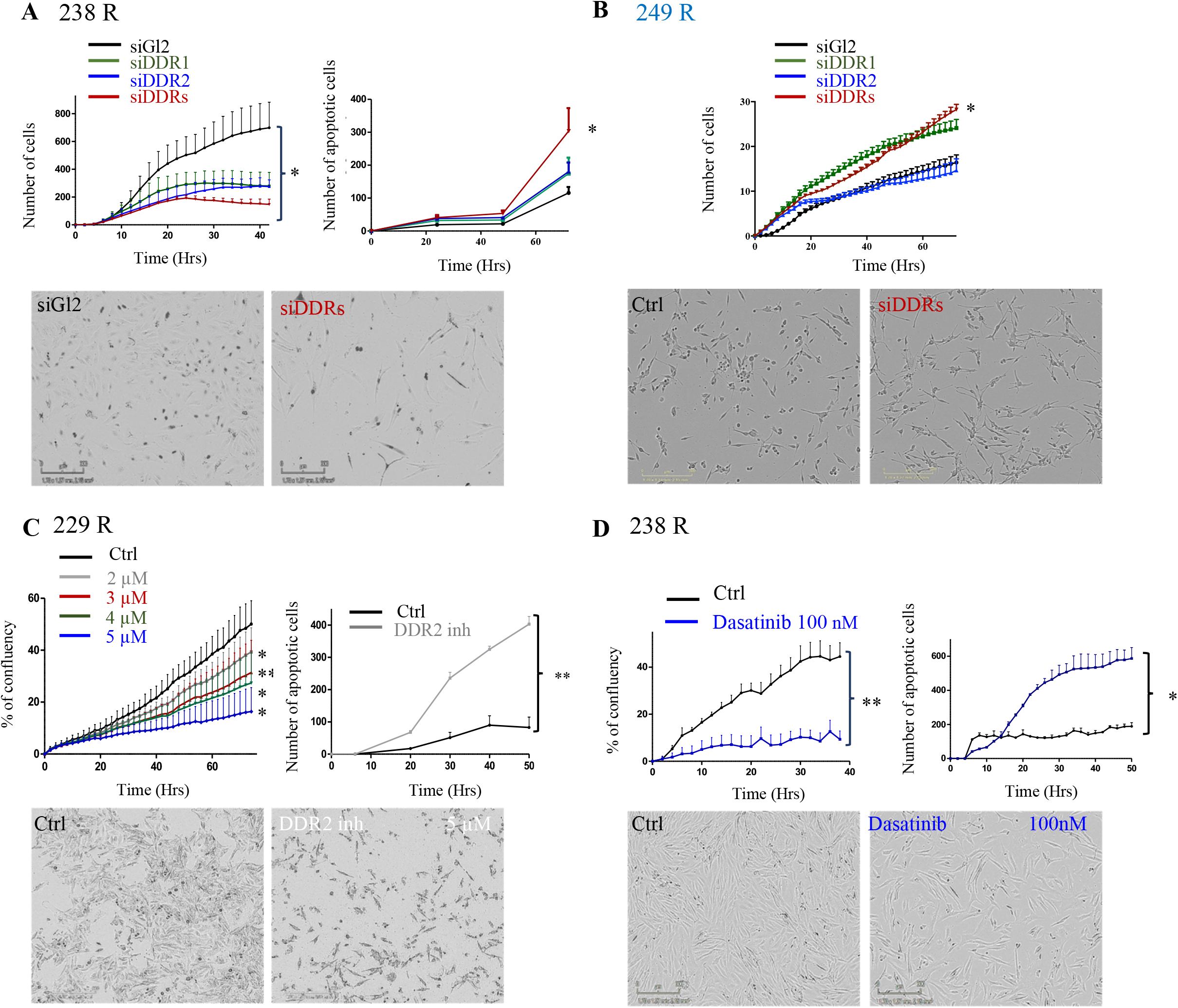
DDRs are involved in resistant tumor cell proliferation. **A**) Left panel: Incucyte® proliferation assay analysis of 238 R cells seeded at 5 000 cells per well in a 96-well plate. The cells were transfected with an siRNA control (siGl2) or targeting DDR1 (siDDR1), DDR2 (siDDR2), or both (siDDR1&2). Values are expressed as the mean ± SEM of three independent experiments. **p<0.01. Right panel: Incucyte® apoptosis assay analysis of 238 R cells seeded at 5 000 cells per well in a 96 well plate. The cells were transfected with an siRNA control (siGl2) or targeting DDR1 (siDDR1), DDR2 (siDDR2), or both (siDDR1&2). Values are expressed as the mean ± SEM of three independent experiments. *p<0.05. **B**) Incucyte® proliferation assay analysis of 249 R cells seeded at 5 000 cells per well in a 96-well plate. The cells were transfected with an siRNA control (siGl2) or targeting DDR1 (siDDR1), DDR2 (siDDR2), or both (siDDR1&2). Values are expressed as the mean ± SEM of three independent experiments. *p<0.05. **C**) Left panel: Incucyte® proliferation assay analysis of 229 R cells seeded at 5 000 cells per well in a 96-well plate cultured in the presence or absence of DDR2 inhibitor. Values are expressed as the mean ± SEM of three independent experiments. *p<0.05, **p<0.01, ***p<0.001. Right panel: Incucyte® apoptosis assay analysis of 229 R cells seeded at 5 000 cells per well in a 96-well plate cultured in the presence or absence of DDR2 inhibitor. Values are expressed as the mean ± SEM of three independent experiments. **p<0.01. **D**) Left panel: Incucyte® proliferation assay analysis of 238 R cells seeded at 5 000 cells per well in a 96-well plate cultured in the presence or absence of 100 nM dasatinib. Values are expressed as the mean ± SEM of three independent experiments. **p<0.01. Right panel: Incucyte® apoptosis assay analysis of 238 R cells seeded at 5 000 cells per well in a 96 well plate cultured in the presence or absence of dasatinib (100 nM). Values are expressed as the mean ± SEM of three independent experiments. *p<0.05.

Treatment of 238 R cells with a DDR1 selective inhibitor decreased DDR1 phosphorylation, inhibited cell proliferation, and inhibited apoptosis, but on the other hand it increased ERK phosphorylation (**Supplemental 3C, supplemental 4B**). However, treatment of resistant melanoma cells with a DDR2 inhibitor decreased both DDR2 and ERK phosphorylation, and consequently inhibited cell proliferation and increased the number of apoptotic cells (**Figure 5C, supplemental 4C**), thus confirming the result obtained in the siRNA experiment.

We also observed an inhibition of tumor cell proliferation and an increase in apoptosis when resistant melanoma cells were treated with dasatinib at 100 nM (concentration at which DDR kinase activity is totally inactivated) compared to cells not treated with dasatinib (**Figure 5D, supplemental 3D & 4D**). Using GSEA on the ratio of siGL2/siDDR2 and control/Dasatinib, we confirmed that cell proliferation and apoptosis signaling pathways were commonly deregulated in these two conditions (**Supplemental 4E)**.

These results indicate that DDRs plays a role in the proliferation exclusively of resistant invasive melanoma cells but not in proliferative resistant melanoma cells.

### The role of DDRs in a physiological 3D model

To analyze the role of DDRs in more physiological conditions, we examined the role of DDRs in cell proliferation using a 3D spheroid culture. The resistant invasive cells seeded in nonadherent conditions had the ability to form spheroids 72 h after seeding (**Figure 6A**), whereas cells not having undergone phenotype switching did not form spheroids. In order to study the impact of DDRs role spheroid maintenance, we treated the spheroids 72h after their formation with DDR1 inhibitor, DDR2 inhibitor or with Dasatinib. Maintenance of the spheroid corresponding to the ability of the cells to form micro-tumors with cell junctions and cell cohesions. Surprisingly, DDR1 inhibition had no impact on spheroid maintenance (**Figure 6B**). We then treated the spheroids 72 h after their formation with the DDR2 inhibitor (5 μM). We found that DDR2 inhibition led to an alteration in spheroid maintenance and a decrease in spheroid area (**Figure 6C**). Together with our previous results showing that resistant cells treated with the DDR2 inhibitor were apoptotic in 2D culture, these results further suggest that the DDR2 inhibitor induces cell death in resistant melanoma cell lines (**Figure 5C, supplemental 4C**). Overall, these results show that DDR2 plays a role in spheroid maintenance.

**Figure 6:**
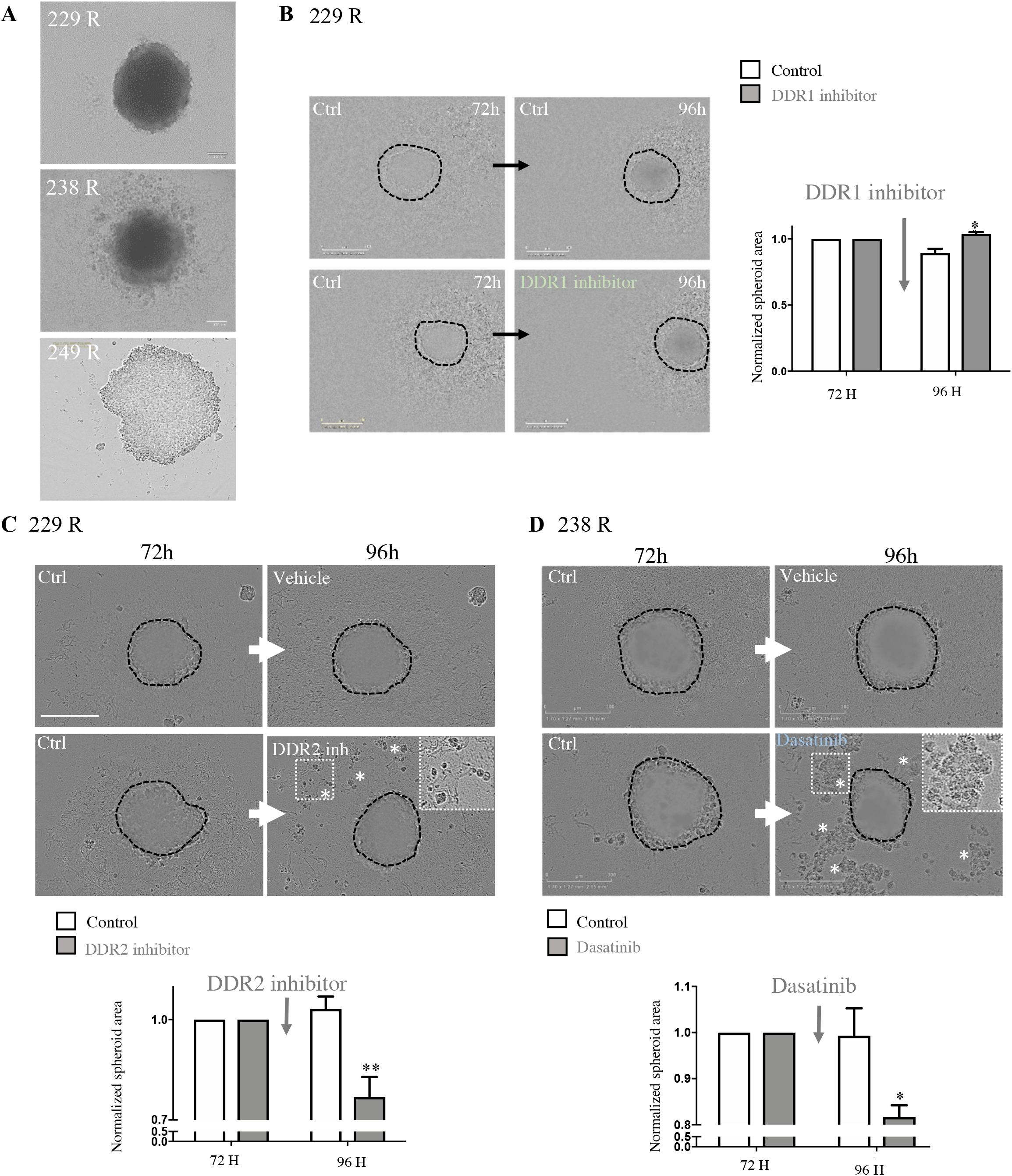
The role of DDRs in a physiological 3D model. **A)** 229R, 238R, and 249R cells were seeded to form spheroids. **B**) 229 R cell spheroids were treated after 72 h with DDR1 inhibitor at 0.8 μM. The graph shows the quantification of spheroid area in the different conditions. Values are expressed as the mean ± SEM of three independent experiments. *p<0.05. **C**) 229 R cell spheroids were treated after 72 h with DDR2 inhibitor at 5 μM. The graph shows the quantification of spheroid area in the different conditions. Values are expressed as the mean ± SEM of three independent experiments. **p<0.01. **D**) 238 R cells were seeded to form spheroids and treated after 72 h with dasatinib at 100 nM. The graph shows the quantification of spheroid area in the different conditions. Values are expressed as the mean ± SEM of three independent experiments. *p<0.05.

We next analyzed the effect of dasatinib on spheroid maintenance by treating spheroids with dasatinib at 100 nM 72 h after seeding. Indeed, we observed a disruption of spheroids in the dasatinib-treated condition (**Figure 6D, supplemental 5**). Spheroid area quantification demonstrated that spheroid area significantly decreased in the dasatinib-treated condition (**Figure 6D, supplemental 5**). We previously showed that cells treated with dasatinib were apoptotic, suggesting that dasatinib induces cell death in resistant melanoma cell lines (**Figure 5D**). These results demonstrated that DDR2 is required for the maintenance of spheroids.

Altogether, these *in vitro* data show that DDR2 is a potential target in resistant melanoma cell proliferation that can be targeted by dasatinib or DDR2 inhibition.

### The role of DDRs in resistant tumor progression *in vivo*

An *in vivo* validation of the effect of dasatinib treatment was necessary to fully confirm the potential of targeting DDRs in vemurafenib-resistant melanoma cells and in turn to counteract this mechanism. Accordingly, we analyzed the effect of dasatinib in a xenograft mouse model of melanoma cell resistance. Firstly, we subcutaneously implanted 229 R cells in immunodeficient NSG mice in order to reproduce the clinical situation of patients harboring resistant metastatic melanoma with the *BRAF* V600E mutation. When tumors reached a volume of 150 mm^3^, the mice were separated into two groups: one group treated with dasatinib by oral gavage, and a control group ongoing treatment with vemurafenib by oral gavage (**Figure 7A**). We observed a stabilization in tumor growth of mice treated with dasatinib compared to the control group, in which tumor growth dramatically increased (**Figure 7B-C**). Western blot analysis of mouse tumors treated with dasatinib showed a decrease in DDR1 and DDR2 phosphorylation compared to a control mouse treated with vemurafenib (**Figure 7D**). These results indicate that dasatinib inhibited both DDR1 and DDR2 phosphorylation *in vivo*. Moreover, we demonstrated that half of the mice treated with vemurafenib presented metastasis to different locations, such as axilla and lymph nodes, whereas mice treated with dasatinib did not display metastasis (**Figure 7E**). We additionally analyzed and compared tissue sections from mice tumors treated with dasatinib or vemurafenib. We observed necrotic areas in dasatinib-treated mice compared to mice treated with vemurafenib (**Figure 7F**). We confirmed this result by localization of Annexin V labeling to the necrotic areas of dasatinib-treated tumors (**Figure 7F**). This result is consistent with our data obtained *in vitro* in both 2D cell and 3D spheroid cultures (**Figures 5&6**).

**Figure 7:**
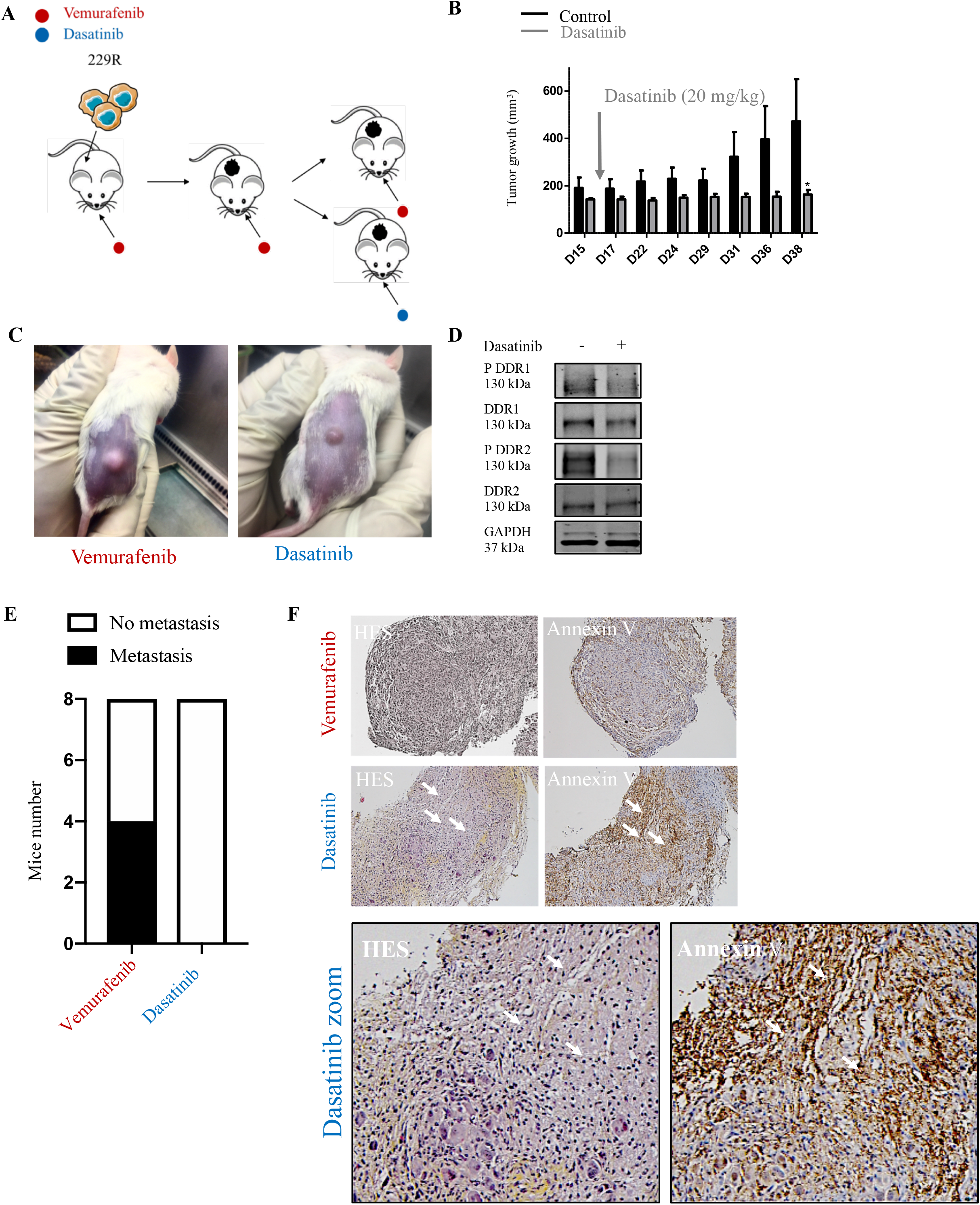
The role of DDRs in resistant tumor progression *in vivo*. **A**) 229 R cells (5*10^6^) were subcutaneously implanted into the right flanks of anesthetized 8-week-old NOD/LtSz-*scid* IL2Rγ*null* (NSG) mice. Mice were treated with vemurafenib until the tumors reached approximately 150 mm^3^ in volume, then the mice were randomly assigned into 2 groups: one control group treated with ongoing treatment with vemurafenib (40 mg/kg), and a second group with mice treated with dasatinib (20 mg/kg) (n=5 in each group). **B**) Tumor growth of 229 R cells in the right flanks of mice. **C**) Photographs of mice treated with vemurafenib or dasatinib. **D**) Western blot analysis of DDR1, DDR2, PDDR1, and PDDR2 expression in primary tumors treated with or without dasatinib. GAPDH was used as the endogenous loading control. **E**) Graphic representation of mice presenting metastasis under vemurafenib or dasatinib treatment. **F**) Immunohistochemistry of primary tumors treated with or without dasatinib. Left panel: HES of primary tumor. Right panel: Immunostaining of Annexin V.

These results demonstrate that dasatinib harbors a pro-apoptotic activity in tumors from subcutaneously implanted vemurafenib-resistant melanoma cells. In conclusion, all these findings highlight that DDR1 and DDR2 could be new therapeutic targets in patients with metastatic melanoma resistant to anti-BRAF therapies.

## Discussion

In 70% of acquired resistance, melanoma cells switch their phenotype and become more aggressive and invasive. Once, this phenotype switching is acquired, resistance to BRAF leads to an over-activation of MAP kinase. It has been demonstrated that DDRs are major players in melanoma. For instance, DDR1 expression is associated with bad prognosis and promotes invasion and migration^44^. Regarding DDR2, it can promote melanoma cell invasion by Jnk activation or proliferation via NFκB/Erk activation^45^.

Firstly, here we demonstrate that expression of DDRs is found in metastatic melanoma, whereas in benign nevi there is no DDR expression. These results are in concordance with *De Moura et al*, showing that DDR1 is associated with bad prognosis in patients with melanoma^44^. Using patient sample data, we highlight that expression of DDR1, DDR2, or both increases after treatment with anti-BRAF.

In the majority of resistant cases, melanoma cells tend to switch their molecular and cellular phenotypes in order to became much more invasive and aggressive. This phenotype switching is characterized by MITF low, AXL high, and actin cytoskeleton remodeling^13–15,46^. Herein, we demonstrate that DDR1 and DDR2 are specifically overexpressed at the protein level in melanoma cells after phenotype switching. These results establish a link between DDR overexpression and phenotype switching. However, the difference between DDR1 protein and mRNA expression levels in the resistant invasive cells could be explained by the fact that DDR1 mRNA expression can be post-transcriptionally regulated for instance by microRNAs^47^. Indeed, microRNA-199a-5p is inversely correlated to the level of expression of DDR1 in ovarian cancer, hepatocellular carcinoma, and colon cancer^47^. In colon cancer, loss of miR-199a-5p induces an up-regulation of DDR1 expression, promoting EMT^48^. Interestingly, a study on melanoma has shown that low expression of miR-199a-5p was associated with bad prognosis and correlated with advanced tumor stage^49^. It could be thus interesting to study a potential regulation of DDR1 by miR-199a-5p in resistant invasive melanoma cells. However, DDR2 overexpression seems more important than that of DDR1 in resistant melanoma cells compared to sensitive cells. This could be due to the fact that resistant invasive melanoma cells secrete their own matrix^50^. Indeed, we demonstrate here that DDR1 expression is reduced when cells are seeded on collagen I. On the other hand, *Sekiya et al* showed in hepatic stellate cells that DDR2 mRNA expression can be decreased by miR-29b, which targets collagen I, suggesting that there is a positive correlation between collagen I expression and DDR2^51^. Accordingly, the tumor microenvironment plays a critical role in vemurafenib-resistant melanoma. Furthermore, DDRs are able to bind to the extracellular matrix. A large number of studies have demonstrated a role of DDRs in cancer cells^26,27^. Nevertheless, DDR1 and DDR2 could be expressed by not only the cancer cells, but also by the stromal cells promoting cell invasion and metastasis^52–54^. Therefore, it will be important to study their expression in melanoma stromal cells.

Contrary to DDR1, little is known about the role of DDR2 in the acquired tumor cell resistance to chemotherapy. Most studies on DDRs in melanoma focus only on one of the two receptors; but none have studied the role of both DDR receptors. Here, we highlight distinct and shared roles of both DDRs in targeted therapy-resistant melanoma.

To begin with, we show that DDR overexpression measured only in invasive resistant cells correlates with AXL expression, a tyrosine kinase receptor which is involved in tumor proliferation, survival, metastasis and resistance to therapy^55^. More precisely, we demonstrate that DDR2 could regulate AXL expression. However, the molecular mechanism by which DDR2 could activate AXL, as well as the signaling pathways induced by this activation, remain unknown. Various studies have reported a role of DDR2 in promoting EMT by SNAIL stabilization in breast cancer or by mTOR activation in gastric cancer^56,57^. Furthermore, Twist stabilizes DDR2 in ovarian cancers promoting EMT^58^. Indeed, given phenotype switching is EMT-like, DDR2 could promote this phenotype switching through regulation of AXL.

Moreover, we demonstrate that DDR2 depletion or inhibition induces a decrease in RhoA signaling by proteomic analysis. These results corroborate a recent study showing an accumulation of actin stress fibers in resistant cells due to RhoA induction, which in turn activates MRTF and YAP^40^. Our proteomic data were confirmed by a decrease in stress fiber formation and RhoA activity when DDR2 was depleted or inhibited. Besides, a DDR2 overexpression in cells not having undergone phenotype switching promoted an increase in stress fiber formation. A role for DDR1 in RhoA activation has already been demonstrated in epidermoid carcinoma cells and the induction of collective migration^59^, and similarly here we report a specific role of DDR2 in RhoA activation promoting stress fiber formation in melanoma. Indeed, it is well known that AXL could activate the RhoA signaling pathway. It will be undeniably important to study if there is any relationship between DDR2 and AXL-promoted RhoA activation and cytoskeletal remodeling^60^.

All these results indicate a tendency for the requirement of DDR2 at the beginning of resistance by promoting phenotype switching via AXL expression and stress fiber formation. It could be relevant next to study if DDR overexpression is sequential during the establishment of resistance in invasive melanoma cells.

In this study, in invasive resistant melanoma cells, we establish that DDR2 depletion or specific inhibition decreases tumor cell proliferation by reducing MAP kinase signaling activity. Likewise, DDR1 depletion or inactivation induces a decrease in cell proliferation, but not through MAP kinase over-activation. Further analysis is required to determine which pathways are activated by DDR1 in targeted therapy-resistant melanoma. To test its clinical relevance, we selected dasatinib, an FDA-approved kinase domain inhibitor used in the treatment of chronic myeloid leukemia^61^. Dasatinib is also known as an inhibitor of SRC, which is involved in a regulatory loop with DDRs^43^. We demonstrate *in vitro* that DDR inactivation by dasatinib inhibits melanoma cell proliferation.

We additionally determine that DDR inactivation induces cell apoptosis (in 2D culture). We show that DDR2 inactivation induces an alteration in spheroid maintenance, whereas DDR1 inhibition shows no effect. It could thus be important to study DDR localization in these conditions given that DDR1, but not yet DDR2, is known to play a role in cell-cell junctions^59^. Here we could postulate that DDR2, but not DDR1, is localized to cell-cell junctions. Indeed, DDRs can be localized to various subcellular compartments associated with several cellular processes involved in cancer progression^27^. For example, there is evidence that DDR1 and DDR2 co-localize along the same collagen I fibers and participate in its degradation^27,62^.

In order to confirm our *in vitro* results, we tested the *in vivo* effects of dasatinib treatment in a xenograft mouse model. We observed that dasatinib treatment blocks proliferation of vemurafenib-resistant melanoma cells. We opted for testing dasatinib treatment because DDRs can be targeted using various FDA-approved tyrosine kinase receptor inhibitors, including dasatinib, as well as imatinib and nilotinib, with IC50s in the low nanomolar range^43^. Nilotinib and dasatinib are more potent inhibitors of DDR1 and DDR2 than imatinib^43^. The IC50 for dasatinib is 0.5 nM for DDR1 and 1.4 nM for DDR2, whereas the IC50 for nilotinib is 43 nM for DDR1 and 55 nM for DDR2^63^. In clinical practice, dasatinib treatment would be more effective than nilotinib due to its high affinity for DDRs and the lower treatment concentration could reduce toxicity. Furthermore, dasatinib has proved efficacious in different cancer types, such as lung, gastric, head, neck, and lung adenocarcinoma^64–67^. Therefore, to abrogate the toxicity due to combination therapy or alternate treatments, sequential treatment is a crucial question in clinical practice^68^. A recent study has demonstrated that switching from MAPK inhibitor therapy to vorinostat (histone deacetylase inhibitor) is more effective in eradicating drug-resistant cells than a drug holiday^68^. In this respect, it would be important to study the effects of long-term sequential treatment in order to delay the onset of resistance to vemurafenib or dasatinib^68^.

Overall, we highlight that DDRs could be good targets in targeted therapy-resistant melanoma and dasatinib can inhibit them. However, dasatinib is not DDR-specific. In recent years, selective DDR inhibitors have emerged. For instance, DDR1 could be targeted specifically by a pyrazolopyramidine alkyne derivative (7rh and 7rj), DDR1-IN-1, or DDR1-IN-2^69,70^. Prof. Longmore’s team have developed selective DDR2 inhibitors, WRG28 and analog CR-13452, that we use in this study. These act via the extracellular domain of the receptor in an allosteric manner^39^. However, the emergence of small molecules and inhibitors that specifically target DDRs could potentially provide progress in cancer treatment development. Indeed, the development of DDR inhibitors capable of targeting both DDRs could be valuable.

To summarize, in this study we uncover shared and distinct roles for DDR1 and DDR2 in targeted therapy-resistant melanoma. We highlight an important role for DDR2 at the beginning of this resistance in melanoma cell phenotype switching. Once an invasive state is acquired, both DDRs promote proliferation of resistant invasive melanoma cells, but only DDR2 is able to over-activate the MAP kinase pathway (**Figure 8A**). Following the perusal of further study, we could expect dasatinib to undergo clinical trials in light of developing a patient management algorithm specifically for those treated with targeted dual therapy-resistance and who are not included in an immunotherapy protocol (**Figure 8B**).

**Figure 8:**
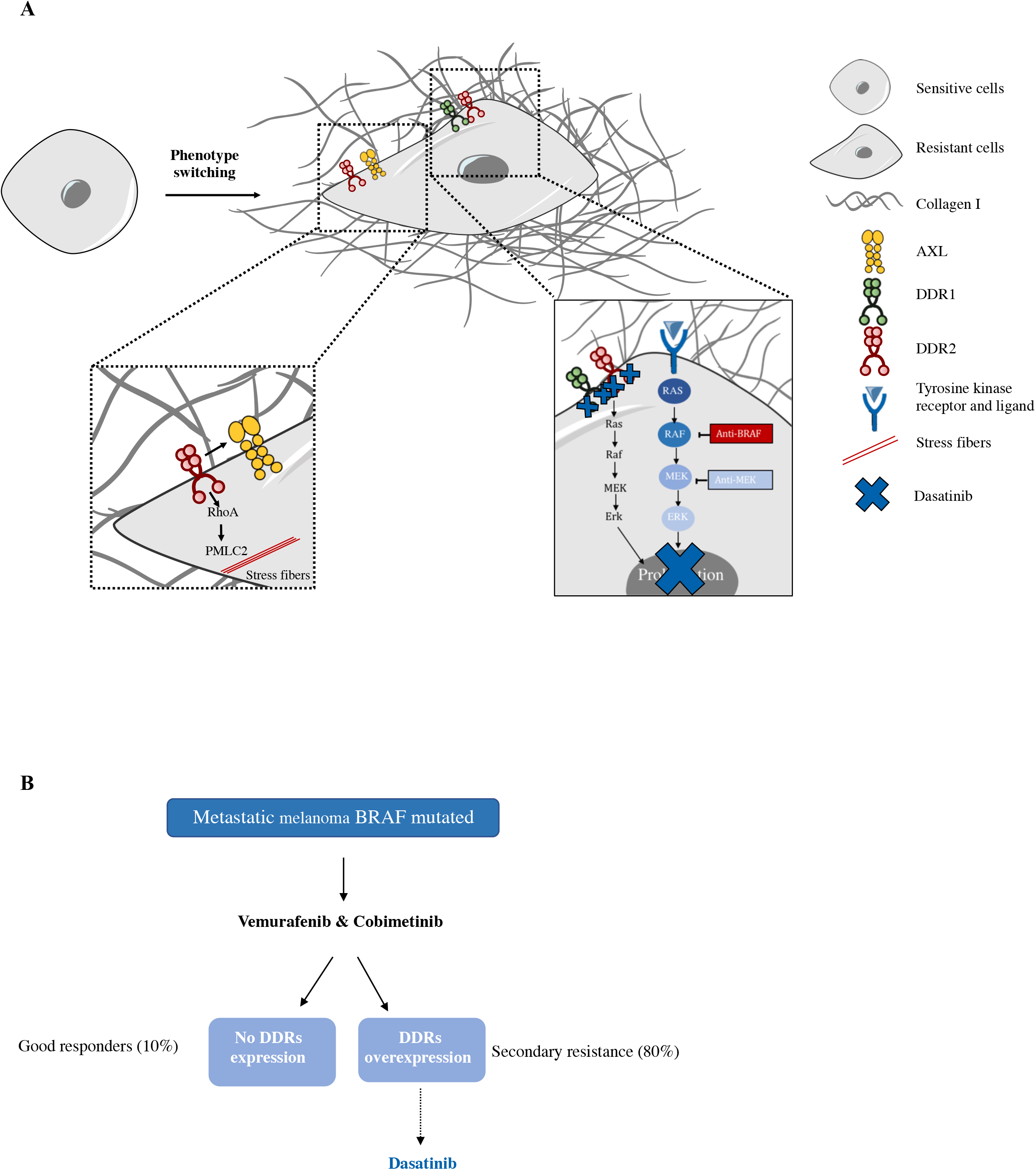
Schematic representation of the distinct roles of DDR1 and DDR2 in vemurafenib-resistant melanoma. **A**) Following BRAF/MEK inhibitor treatment, BRAF-mutant melanoma cell lines undergo phenotype switching and became invasive via DDR2 upregulation. This in turn could i) regulate AXL expression and ii) activate the RhoA signaling pathway promoting cytoskeletal modifications. Once this resistant invasive phenotype is acquired, melanoma cells require upregulation of DDR1 and DDR2 to promote proliferation, but only DDR2 is able to overactivate the MAP kinase pathway. **B**) Proposed model for the treatment of patients with BRAF-mutant metastatic melanoma. For vemurafenib-resistant patients, we propose post-treatment biopsy analysis. If DDR expression is detected at similar levels as to before treatment, we put forward dasatinib as an alternative treatment.

## Declaration

### Ethics approval and consent to participate

The institutional animal ethics committee of Bordeaux University approved all animal use procedures and all efforts were made to minimize animal suffering.

## Acknowledgements

We are grateful to Prof. Longmore for providing us with DDR2 inhibitors. Many thanks to Dr David Santamaria for giving us advice on this project and for providing us with a list of MAP kinase targets and their associated primers. We are also grateful to Dr Sophie Tartare-Deckert for providing us with melanoma cells. We are grateful to the Centre de Ressources Biologiques (CRB), Bordeaux, for access to melanoma patient samples.

## Supplemental figures

**Supplemental 1:**
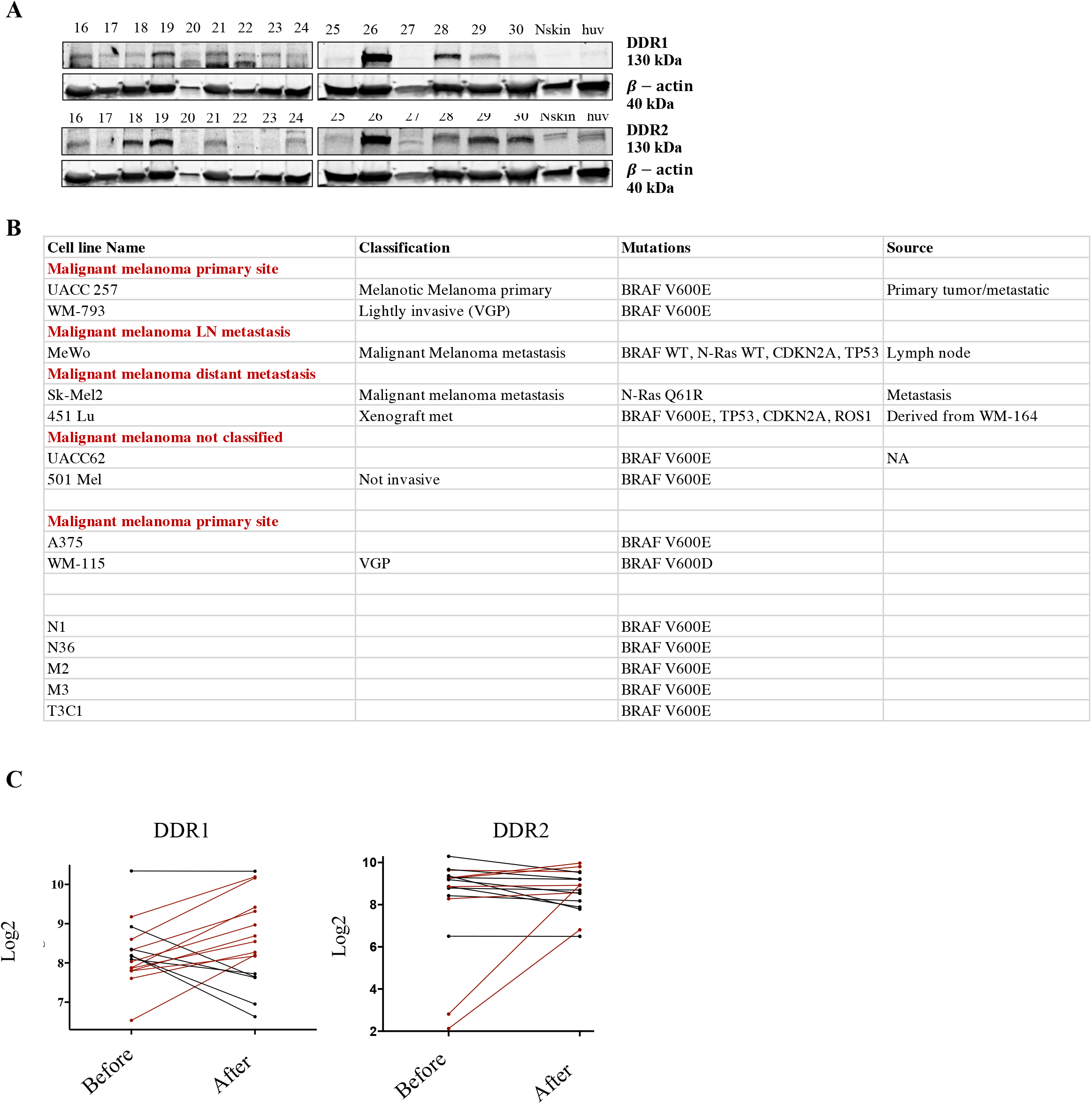
DDR expression in melanoma and in targeted therapy-resistant melanoma cells. **A**) Western blot analysis of DDR1 and DDR2 expression in 15 metastatic melanoma patient biopsies. β-actin was used as the endogenous loading control. **B)** Summary table of the different cell lines used in the screening experiment as well as the mutations found in these cell lines. **C**) RNA sequencing data (*GSE50509*) for DDR1 or DDR2 mRNA expression before and after treatment with an anti-BRAF.

**Supplemental 2:**
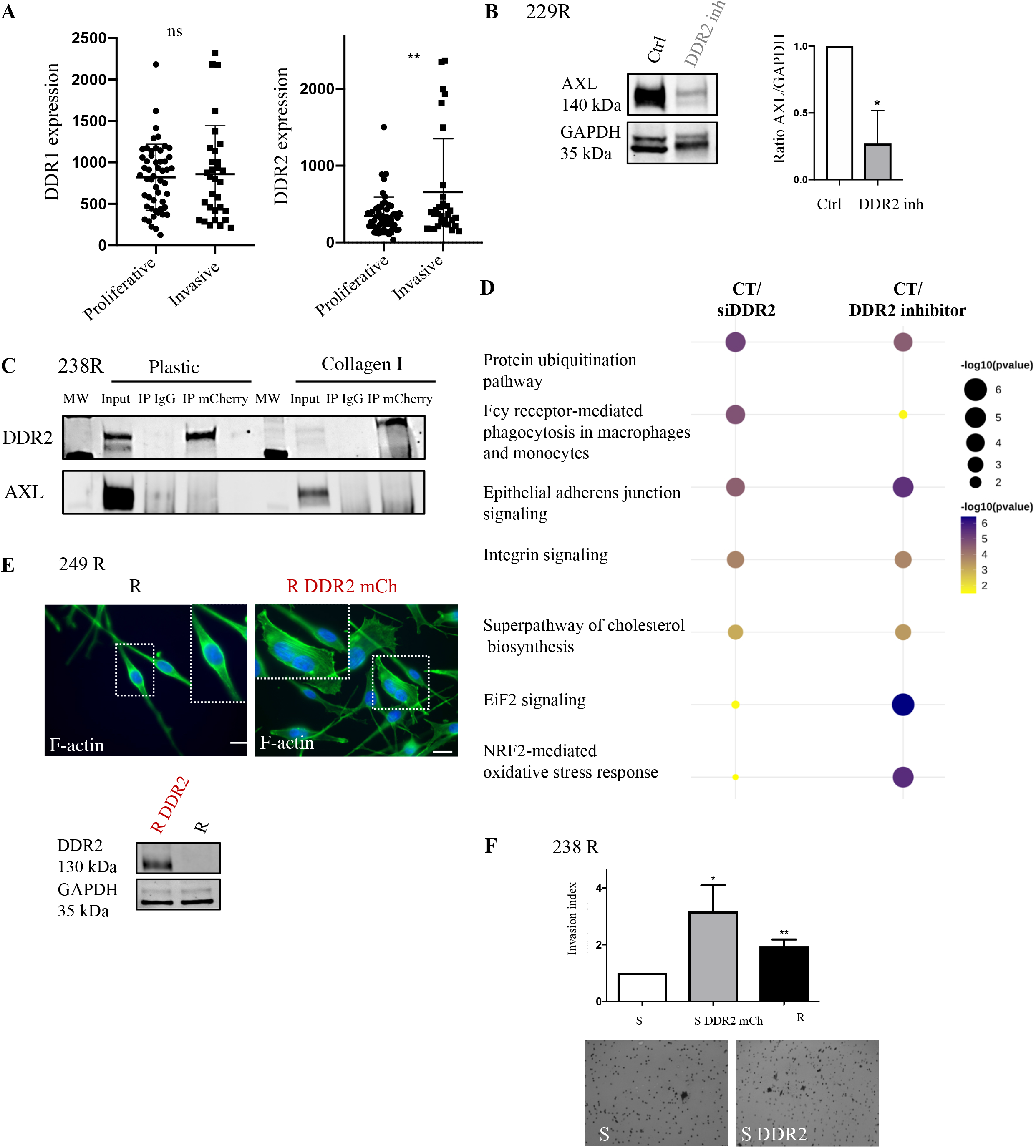
DDR2 involvement in phenotype switching. **A**) RNA sequencing data (GSE4845, GSE4843, GSE4840) for DDR1 and DDR2 mRNA expression in proliferative or invasive melanoma cells. **B**) 229 R cells were treated with DDR2 inhibitor. Protein extracts were then analyzed by immunoblotting to determine PDDR2, DDR2, and AXL expression. GAPDH was used as the endogenous loading control. The graph shows the quantification of AXL/GAPDH expression. Values are expressed as the mean ± SEM of three independent experiments. *p<0.05. **C**) Study of the interaction between AXL and DDR2 by co-immunoprecipitation (IP) assay in 238 R cells. AXL or DDR2 were immunoprecipitated using AXL or mCherry antibodies. **D**) Other pathways commonly enriched in the following conditions: 238 R cells transfected with siRNA targeting DDR2 or treated with DDR2 inhibitor. Gene set enrichment analysis (GSEA) was performed against the Ingenuity Pathways database (Fisher’s Exact test expressed in −log10pvalue). **E**) 249 R cells were transiently transfected with DDR2-mCherry. Representative images of the actin cytoskeletons of 249 R DDR2 mCh cells. Fluorescent staining corresponds to F-actin (green) and nuclei (blue). Scale bar: 6.78 uM. Protein extracts were then analyzed by immunoblotting to determine DDR2 expression. GAPDH was used as the endogenous loading control. **F**) 238 S cells were transiently transfected with DDR2-mCherry. 238 (S, S DDR2-mCh, R) cells were seeded in Matrigel-coated chambers and invasion was assessed. The graph shows invasion index quantification. Values are expressed as the mean ± SEM of three independent experiments. *p = 0.0221; **p = 0.0057.

**Supplemental 3:**
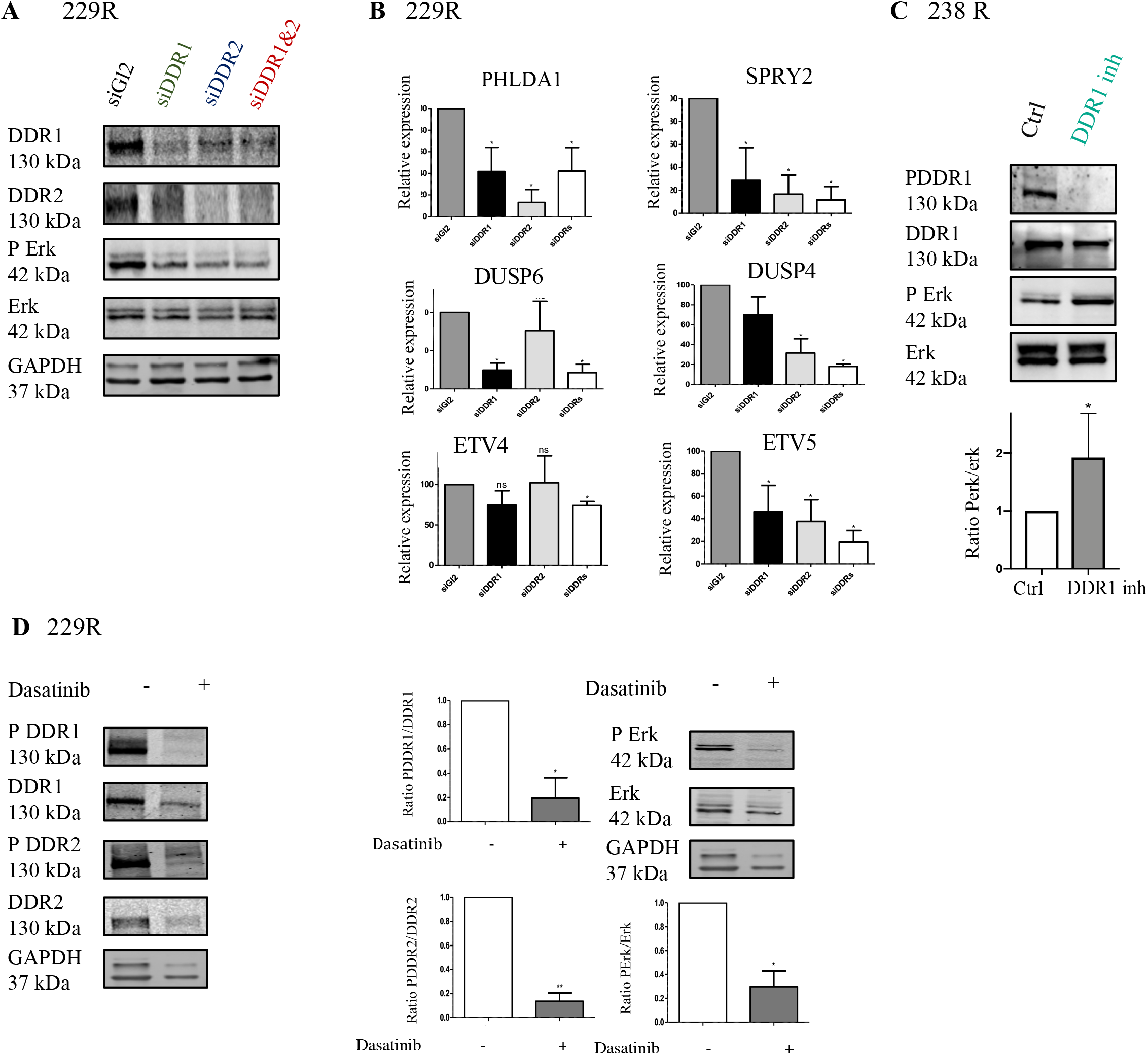
DDR-associated phenotype switching is involved in MAP kinase pathway activation. A) 229 R cells were transfected with an siRNA control (siGl2) or targeting DDR1 (siDDR1), DDR2 (siDDR2), or both (siDDR1&2). Protein extracts were then analyzed by immunoblotting to determine PErk and Erk expression. GAPDH was used as the endogenous loading control. **B**) mRNA expression of MAP kinase targets in 229 R cells. The graph shows quantification of PHLDA1, SPRY2, DUSP6, DUSP4, ETV4, and ETV5 expression. Values are expressed as the mean ± SEM of three independent experiments. *p<0.05. **C**) 238 R cells were treated with DDR1 inhibitor (7rh) for 3 days. Protein extracts were then analyzed by immunoblotting to determine PErk, Erk, PDDR1, and DDR1 expression. The graph shows the quantification of the ratio of PDDR1/DDR1 and PErk/Erk expression. Values are expressed as the mean ± SEM of three independent experiments. *p<0.05. **D**) 229 R cells were treated with dasatinib for 2 hours. Protein extracts were then analyzed by immunoblotting to determine PErk and Erk expression. GAPDH was used as the endogenous loading control. The graph shows the quantification of the ratio of PDDR1/DDR1, PDDR2/DDR2, and PErk/Erk expression. Values are expressed as the mean ± SEM of three independent experiments. *p<0.05, **p<0.01.

**Supplemental figure 4:**
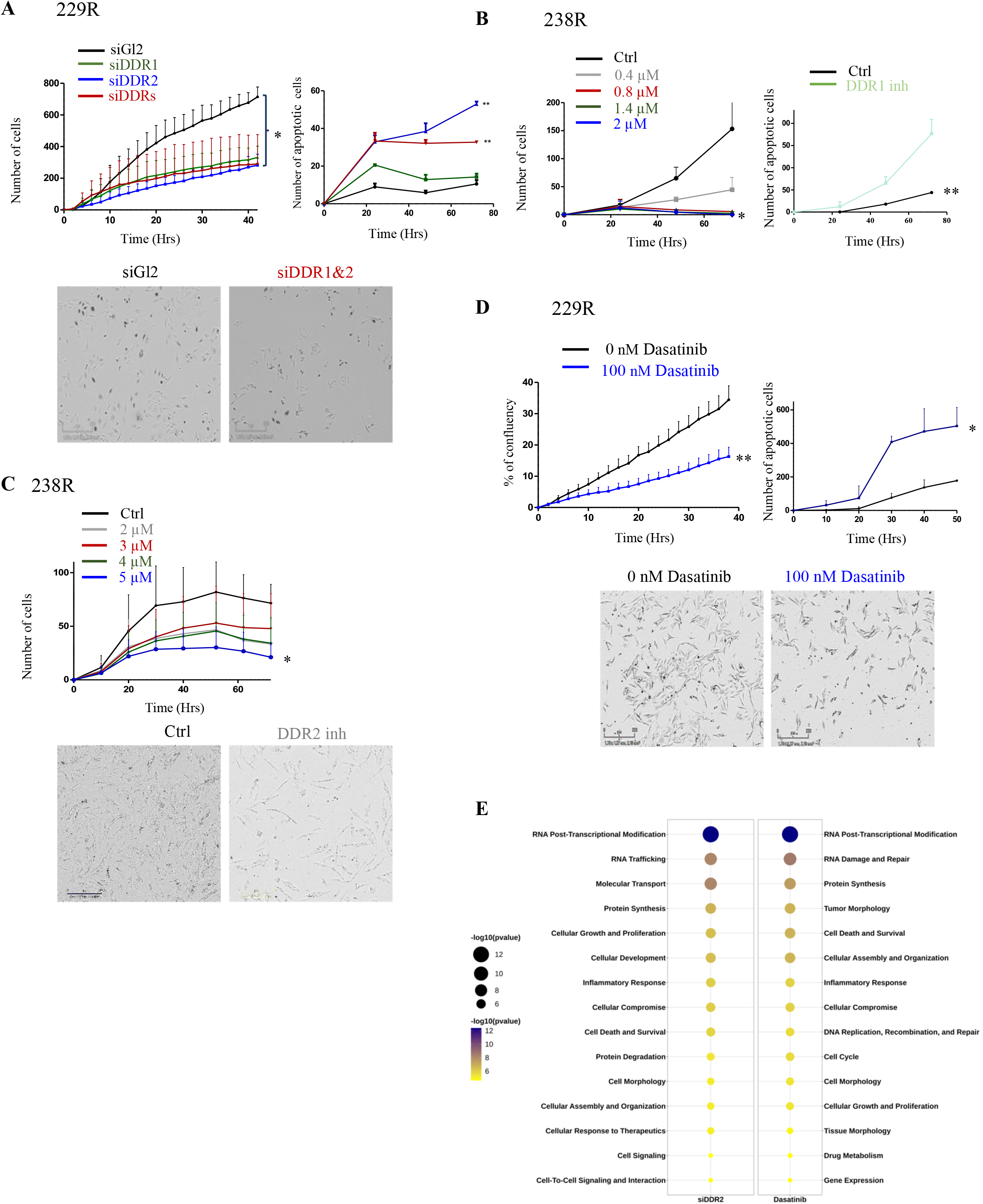
DDRs are involved in resistant tumor cell proliferation. **A**) Left panel: Incucyte^®^ proliferation assay analysis of 229 R cells seeded at 5 000 cells per well in a 96-well plate. The cells were transfected with an siRNA control (siGl2) or targeting DDR1 (siDDR1), DDR2 (siDDR2), or both (siDDR1&2). *p<0.05. Right panel: Incucyte^®^ apoptosis assay of 229 R cells seeded at 5 000 cells per well in a 96-well plate. The cells were transfected with an siRNA control (siGl2) or targeting DDR1 (siDDR1), DDR2 (siDDR2), or both (siDDR1&2). Values are expressed as the mean ± SEM of three independent experiments. *p<0.05. **B**) Left panel: Incucyte® proliferation assay analysis of 238 R cells seeded at 5 000 cells per well in a 96-well plate and cultured in the presence or absence of DDR1 inhibitor. Values are expressed as the mean ± SEM of three independent experiments. *p<0.05. Right panel: Incucyte® apoptosis assay analysis of 238 R cells seeded at 5 000 cells per well in a 96-well plate and cultured in the presence or absence of DDR1 inhibitor. Values are expressed as the mean ± SEM of three independent experiments. **p<0.01. **C**) Incucyte® proliferation assay analysis of 238 R cells seeded at 5 000 cells per well in a 96-well plate and cultured in the presence or absence of DDR2 inhibitor. Values are expressed as the mean ± SEM of three independent experiments. *p<0.05. **D**) Left panel: Incucyte® proliferation assay analysis of 229 R cells seeded at 5 000 cells per well in a 96-well plate and cultured in the presence or absence of 100 nM dasatinib. Values are expressed as the mean ± SEM of three independent experiments. **p<0.01. Lower panel: Incucyte® apoptosis assay analysis of 229 R cells seeded at 5 000 cells per well in a 96-well plate and cultured in the presence or absence of dasatinib (100 nM). Values are expressed as the mean ± SEM of three independent experiments. *p<0.05. **E)** Bubble plot of the pathways commonly and significantly enriched between the following conditions: 238 R cells transfected with siRNA targeting DDR2 or treated with dasatinib. The bubble plot represents the ratio of control/siDDR2 or control/dasatinib. Gene set enrichment analysis (GSEA) was performed against the Ingenuity Pathways database (Fisher’s Exact test expressed in −log10pvalue). Colors and dot size represent minus logarithms of adjusted p-value (padj). Column height represents numbers of genes enriched in a pathway.

**Supplemental figure 5:**
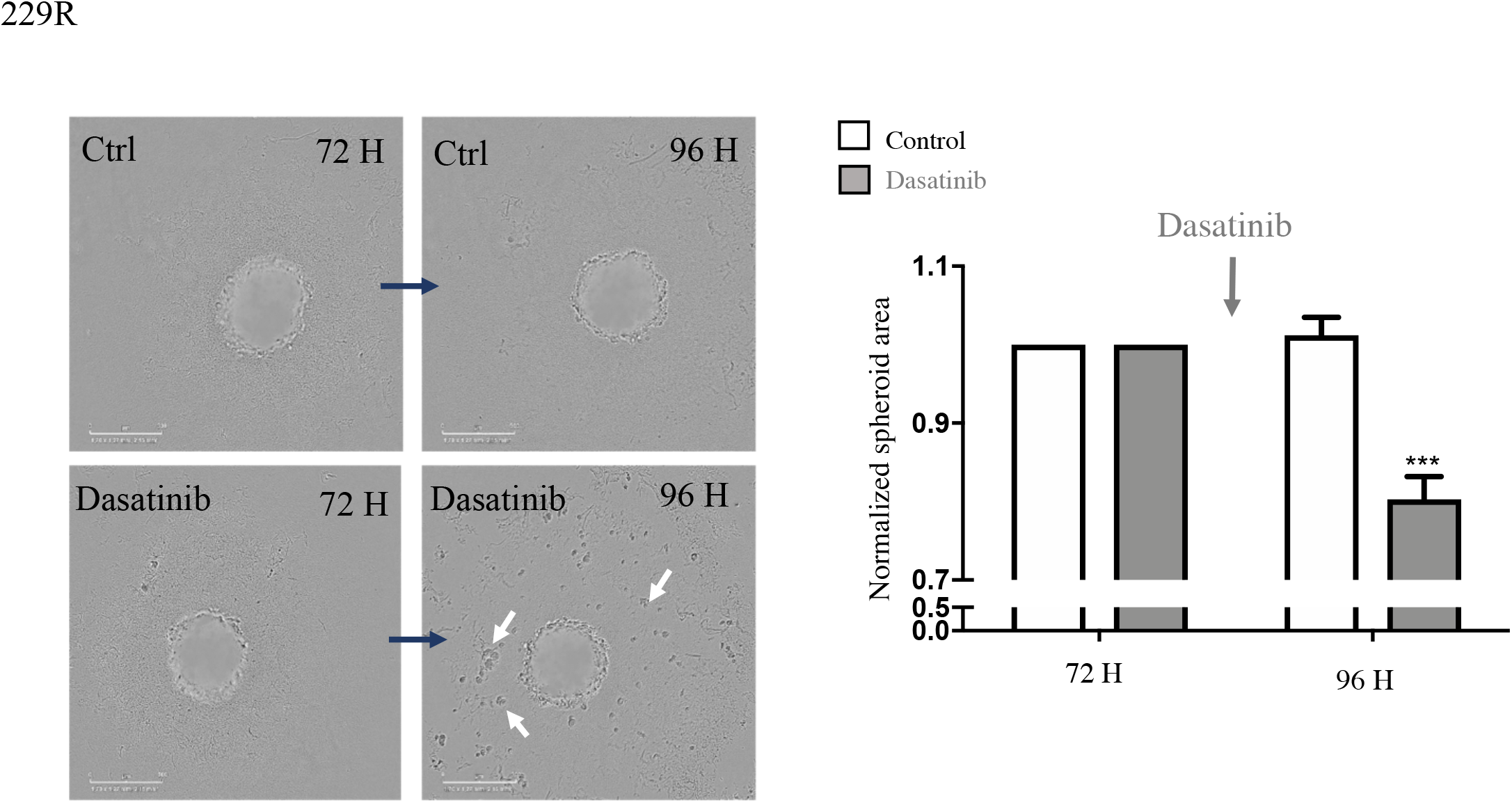
The role of DDRs in a physiological 3D model. 229 R cells were seeded to form spheroids and were treated with dasatinib at 100 nM after 72 h. The graph shows the quantification of spheroid area in the different conditions. Values are expressed as the mean ± SEM of three independent experiments. ***p<0.001.

**Supplemental table 1:**
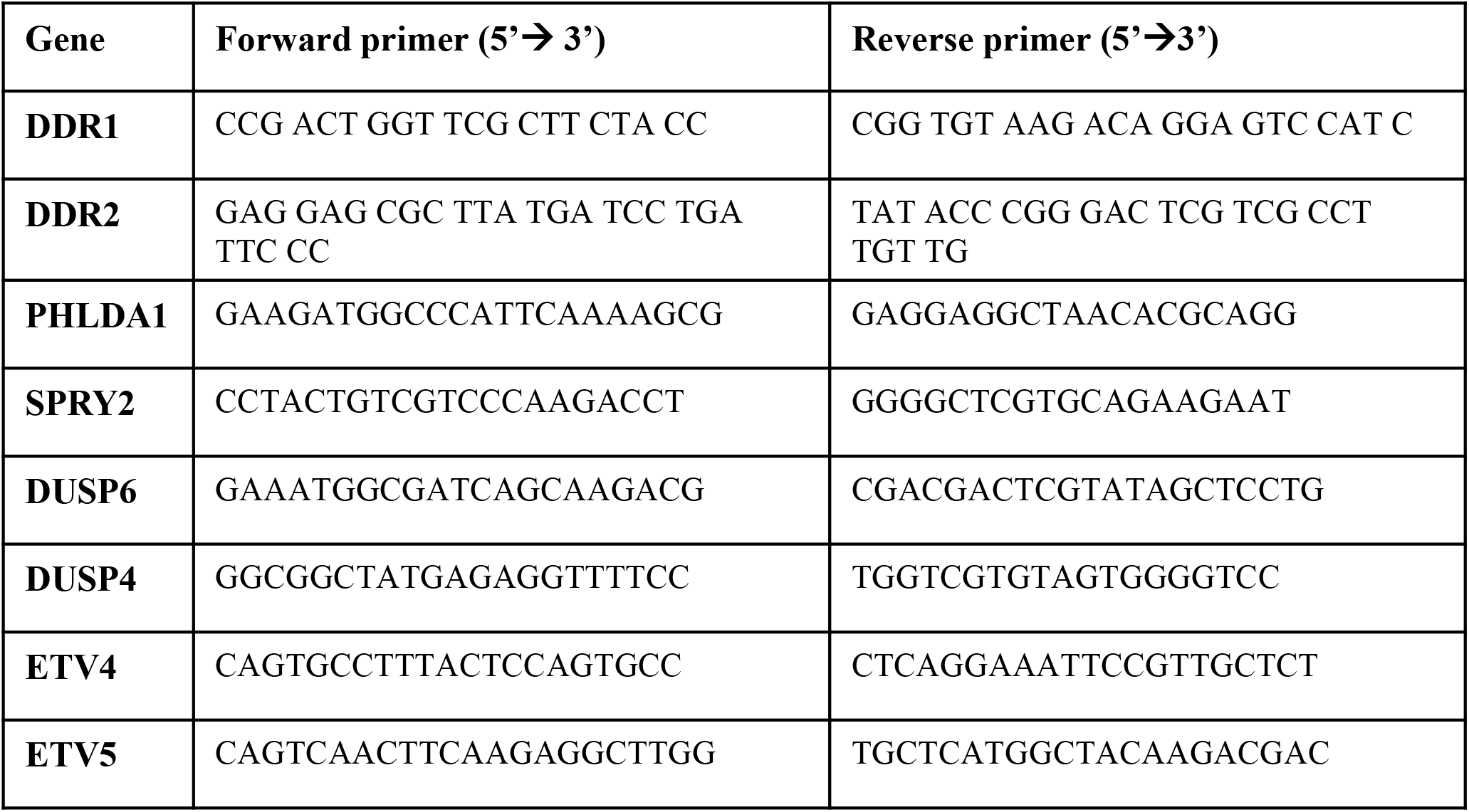
List of PCR primers.

